# Introgression of a dominant phototropin1 mutant superenhances carotenoids and boosts flavor-related volatiles in genome-edited tomato *RIN* mutants

**DOI:** 10.1101/2023.05.05.539534

**Authors:** Narasimha Rao Nizampatnam, Kapil Sharma, Prateek Gupta, Injangbuanang Pamei, Supriya Sarma, Yellamaraju Sreelakshmi, Rameshwar Sharma

## Abstract

The tomato (*Solanum lycopersicum*) ripening inhibitor (*rin*) mutation is known to completely repress fruit ripening. The heterozygous (*RIN*/*rin*) fruits have extended shelf life, ripen normally, but have poor taste and flavour. Even the CRISPR/Cas9-generated *rin* alleles have these undesirable attributes associated with the *rin* mutation. To address this, we used genome editing to generate newer alleles of *RIN* (*rin^CR^*) by targeting the K domain, which is essential for the oligomerization of MADS-box transcription factors. Unlike previously reported CRISPR alleles, the *rin^CR^*alleles displayed delayed onset of ripening, suggesting that the mutated K domain represses the onset of ripening. The *rin^CR^* fruits had extended shelf life and accumulated carotenoids at an intermediate level between *rin* and wild-type parent. Besides, the metabolites and hormonal levels in *rin^CR^* fruits were more akin to *rin*. To overcome the negative attributes of *rin*, we crossed the *rin^CR^* alleles with *Nps1*, which enhances carotenoid levels in tomato fruits. *Nps1* harbours a dominant-negative mutation in the plant photoreceptor phototropin1. The resulting *Nps1*/*rin^CR^* hybrids had extended shelf life and 4.4-7.1-fold higher carotenoid levels than the wild-type parent. The *Nps1*/*rin^CR^* fruits had higher auxin and reduced ABA levels, which are reportedly linked with slower ripening. The metabolome of *Nps1*/*rin^CR^*fruits revealed higher sucrose, malate, and volatiles associated with tomato taste and flavour. Notably, the boosted volatile levels in *Nps1*/*rinCR* were only observed in fruits bearing the homozygous *Nps1* (*Nps1*/*Nps1*) mutation. Our findings suggest that the *Nps1* introgression into tomato ripening mutants provides a promising strategy for developing tomato cultivars with extended shelf life, improved taste, and flavour.

## Introduction

Fruit ripening is a complex process involving a range of physiological and biochemical changes. The ripening initiates the change in the fruit coloration, development of aroma, and softening of fruit flesh. The ripening makes fruit attractive to frugivores and helps in the wide dispersion of seeds. Based on the surge in respiration at the onset of ripening, fruits have been classified as climacteric (banana, melon, and tomato) and nonclimacteric (grape, strawberry, and citrus). Climacteric fruits, like tomatoes, show a significant increase in ethylene emission, which marks the onset of ripening. The tomato (*Solanum lycopersicum*) has emerged as a model system for deciphering the mechanisms underlying ripening due to its commercial importance, well-characterized genome, and genetics, and the availability of several monogenic mutations affecting various aspects of ripening (**Klee and Giovannoni, 2011**). Post-anthesis, tomato fruit grows in size and stores nutrients; the ripening process is suppressed until it reaches maturity. On reaching maturity, a series of highly coordinated physiological and biochemical changes transform the fruit into a palatable and nutritious product.

The knowledge about molecular mechanisms regulating fruit ripening is essential to enhance the nutrients, organoleptic fruit quality, and shelf life and minimize post-harvest losses. The strategies for improving tomato fruit quality have heavily relied on the use of spontaneous mutants, such as the increase in all pigments (*hp-1, hp-2, pd*), carotenoids (*r, t, at, B, Bmo_B_*), chlorophyll (*gf*), flavonoids (*y*) and ripening process (*Nr, rin, Gr*) (**Dono et al., 2020**). Among the mutants regulating the ripening, *ripening inhibitor* (*rin*), *nonripening* (*nor*), and *Colorless nonripening* (*Cnr*) have been extensively used due to their nonripening phenotypes (**Robinson and Tomes, 1968; Tigchelaar et al., 1973; Thompson et al., 1999; Giovannoni, 2004; Manning et al., 2006; Kumar et al., 2018**). The *RIN*, *CNR* (an epigenetic mutation in promotor), and *NOR* genes encode transcription factors belonging to MADS, SBP box, and NAC family, respectively (**Seymour et al., 2013**). These mutations lead to fruits that do not undergo the normal hallmarks of ripening, such as softening, colour change, and sugar accumulation.

Among the non-ripening mutants, *rin* has been extensively studied. It shows a severe ripening-defective phenotype, displaying a lack of fruit softening and an absence of red coloration imparted by lycopene. The *rin* also lacks the climacteric rise of respiration and ethylene, and the fruits do not respond to exogenous ethylene (**Tigchelaar et al., 1978**). The studies using chromatin immunoprecipitation (ChIP) revealed that RIN directly binds to the promotor of at least 241 genes involved in ethylene synthesis/signalling, cell wall modification, carotenoid biosynthesis, and ripening-related TFs such as *NOR*, and *CNR* (**Fujisawa et al., 2013**). The RIN acts as a transcriptional activator and forms complexes with other MADS-box TFs regulating ripening, such as TAGL1, FUL1, and FUL2 (**Fujisawa et al., 2014**). Based on the wide-ranging action of RIN on ripening processes, it was proposed that RIN acts as a master regulator of tomato ripening upstream of ethylene.

The recent studies on CRISPR/Cas9 generated *RIN* mutants revealed inconsistencies between the phenotypes of *rin* and genome-edited *RIN*-knock-out mutant lines. Unlike the *rin* mutant, the genome-edited *RIN* null mutants show partial induction of ripening (**Ito et al., 2015**). The genome editing or RNA interference of the *rin* mutant allele partially reversed the nonripening phenotype of the mutant (**Ito et al., 2017; Li et al., 2018**). These studies led to a revision of the role of *RIN* in tomato ripening. It is now believed that RIN is not necessary for ripening initiation, and the mutant’s severe nonripening phenotype results from the RIN protein’s repressor activity. The *rin* mutant has a large deletion between it and the following gene, *macrocalyx* (*MC*). The deletion led to the generation of an in-frame fusion mRNA that produces a hybrid transcription factor, RIN-MC, which conferred the gain of dominant repressor activity rather than the previously presumed loss of function (**Ito et al., 2017)**.

The comparison of CRISPR-generated alleles of the *RIN* gene revealed the role of different domains in regulating ripening. The ripening phenotype of mutants generated by CRISPR revealed that the loss of gene function does not arrest the ripening initiation; it rather slows the progression and completion of ripening (**Ito et al., 2017**). Since no RIN protein was detected in the above mutants using a MADs domain-specific antibody (70-93 AA), these mutants were considered knockout alleles (*rin-KO*). Likewise, *RIN-CRISPR* alleles encoding for a truncated RIN protein (1-∼78 AA) initiated partial ripening almost at a similar time to wild-type fruits (**Li et al., 2020**). The gene-edited *RIN* alleles accumulated lycopene, and emitted ethylene, albeit at a highly reduced level (**Li et al., 2020; Ito et al., 2017)**. The *RIN-CRISPR* alleles lacked system II ethylene emission, and exogenous propylene did not fully stimulate system II ethylene emission (**Li et al., 2020).** The post-harvest shelf life of these gene-edited mutants did not differ from the wild type, though the mutants showed subdued expression of cell-wall modifying genes (**Li et al., 2020; Ito et al., 2020**).

Another gene-edited *RIN* allele, *rinG2*, partially truncated in the C domain of RIN (199 aa), exhibited a ripening phenotype distinct from the null mutants. The *rinG2* mutant fruits showed extended shelf life and reduced softening rate like the *rin* mutant (**Ito et al., 2020**). Unlike knockout mutants, the immunoblots of *rinG2* showed a truncated RIN protein (**Ito et al., 2021**). Though the *rinG2* allele had extended fruit shelf life, it had negative attributes of the *rin* mutant and had reduced lycopene levels similar to the knockout mutants (**Ito et al., 2020**). An attempt was made to increase the lycopene level in *rinG2* fruits by knocking out the lycopene β-cyclase (*CYCB*) gene. However, homozygous *CYCB* knockouts were lethal, so in *rinG2,* only heterozygous *CYCB* knockouts could be raised, which had moderately increased lycopene levels (**Ito et al., 2020**). In contrast, the crossing of *rinG2* with the wild-type parent restored the lycopene nearly to the wild-type levels while maintaining the extended shelf life of the fruits (**Ito et al., 2021**).

Tomato breeders have utilized the non-ripening characteristic of the *rin* mutant to produce tomato hybrids with an extended shelf life. However, these hybrids have had drawbacks, such as lower levels of lycopene, decreased organoleptic quality, and a loss of the distinctive tomato flavour (**Baldwin et al., 1995**). The loss of flavour and taste in *rin* hybrids was due to the negative impact of *rin* mutation on the formation of ripening-associated metabolites and volatiles (**Osorio et al., 2011, 2020; Wang et al., 2020).** Even in gene-edited *RIN-CRISPR* alleles, the formation of volatiles associated with consumer preferences was adversely affected (**Li et al., 2020**).

The restoration of primary metabolites and aroma-related volatiles is an important target in tomato fruit biology; as compared to heirloom tomatoes, the modern tomato cultivars are poor in taste and flavour (**Tieman et al., 2017**). In this study, we utilized CRISPR to generate four new *RIN* alleles in the K domain; two had in-frame amino acid deletion(s), and the other had protein truncation. Although these alleles exhibited a reduction of lycopene similar to the other gene-edited mutants, the fruits had an extended on-vine and post-harvest shelf life. We overcame the negative traits of taste and flavour associated with *rin* and knockout mutants by crossing our gene-edited *RIN* alleles with a phototropin1 mutant *Nps1.* The *Nps1* mutant dominantly upregulates carotenoids and volatiles in tomatoes (**Kilambi et al., 2021**). We show that the *Nps1/rin^CR^*double hybrids have > 4-fold higher lycopene levels than their wild-type parent, and fruits are enriched in metabolites and volatiles related to consumer preference.

## Results

The mutagenesis of tomato *rin* mutant and *RIN* gene using CRISPR/Cas9 has relegated the function of RIN from an initiator of fruit ripening to the one that prevents overripening (**Ito et al., 2017, 2020; Li et al., 2020**). The restoration of full ripening and augmentation of carotenoids and flavour levels in *RIN* gene-edited alleles have not been investigated. In this study, we investigated whether another mutant could mitigate the negative attributes of *RIN* gene-edited alleles on lycopene and flavour levels.

### Generation of gene-edited *rin* lines

The RIN is a type II MADS domain (IPR2100, https://www.ebi.ac.uk/interpro/entry/InterPro/IPR002100/) protein with a conserved K domain (84-172AA) (IPR002487, https://www.ebi.ac.uk/interpro/entry/InterPro/IPR002487/) that is critical for the MADS protein’s oligomerization (**Lai et al., 2019**). We targeted the K domain for gene knockout using a single guide RNA (sgRNA) (**Figure S1A**). The sgRNA editing of *RIN* disrupted a TaiI restriction enzyme (RE) site (ACGT^), enabling the identification of edited plants by loss of the RE site. The construct used for transformation in pCAMBIA binary vector 2300 backbone had CAS9 driven by 2X35S CaMV promoter and sgRNA driven by U6 promoter (**Nekrasov et al., 2013**) (**Figure S1B**). Of 52 independent T_0_ transgenic lines, 39 were CAS9 positive, and 34 were resistant to TaiI digestion in the PCR-RE method (**Figure S1C**). Out of 34 lines, we randomly picked 14 TaiI*-*resistant T_0_ lines (**Table S1**). From the above lines, four homozygous alleles (*rin^CR1-4^*) were selected in T_1_, and CAS9-free plants were selected in T_2_ generation onwards at the seedling stage (**Figure S2, Dataset S1**).

The *rin^CR1^, rin^CR2^,* and *rin^CR3^*had 3, 6, and 5 base pair deletions, respectively, while *rin^CR4^*had a single base insertion in the gene (**Figure 1A**). Sequencing the predicted top four off-targets in the T_1_ generation revealed no off-target mutations in any of the tested lines, including four *rin^CR^*alleles (**Table S2**). In *rin^CR1^*, *rin^CR2^*base pair deletions led to in-frame amino acid deletion(s); thereby, these lines retained the K domain but with 1 and 2 AA deletion(s), respectively. In *rin^CR3^*and *rin^CR4^*, frameshift mutation created a premature stop codon resulting in protein truncation. Nevertheless, *rin^CR3^* and *rin^CR4^* retained nearly half of the K domain (**Figure 1B**). The CAS9-free *rin^CR1-4^* alleles were then analysed in T_2_ and later generations.

**Figure 1.**
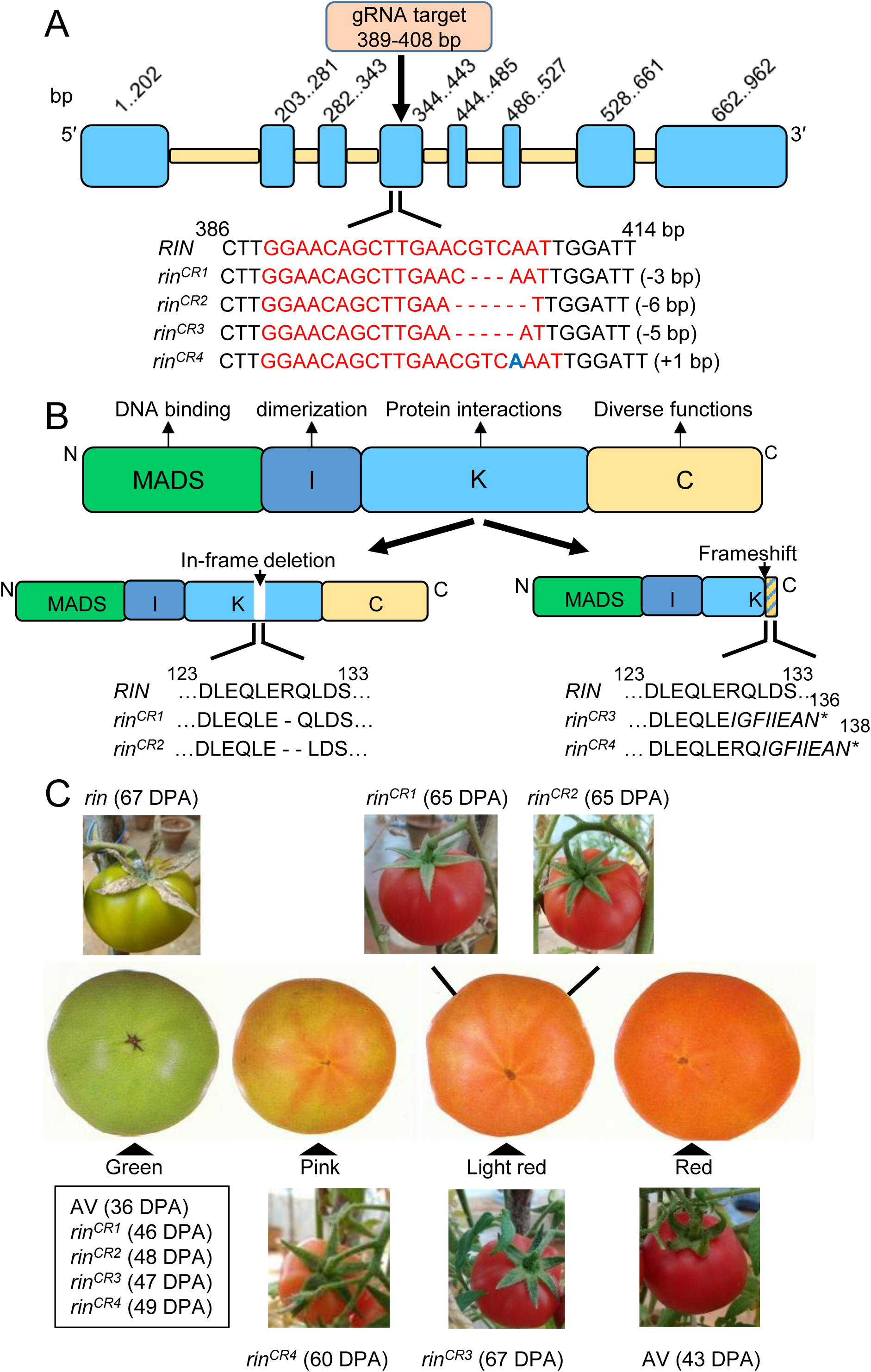
Characterization of genome-edited *rin* alleles. **A.** The exon 4 of the *RIN* gene was edited using a single gRNA, generating four *rin^CR^* alleles. The gRNA target sequence is highlighted in red. The deleted nucleotides are indicated by a dash (-). The insertion of a single nucleotide in the *rin^CR4^* allele is marked in blue. **B.** The cartoon shows four functional domains of the RIN protein. The editing disrupted K-domain with an in-frame deletion of the amino acid(s) in *rin^CR1^* and *rin^CR2^* and frameshift in *rin^CR3^* and *rin^CR4^*, leading to protein truncation. **C.** The progression of ripening in *rin* mutant and *rin^CR^* alleles. Unlike *rin* mutant, which does not ripen, *rin^CR^* alleles show ripening, albeit at a sluggish pace than Arka Vikas (AV). Note, at the final ripe stage, the fruits of *rin^CR^* alleles do not attain deep red coloration like AV. The box below the green stage shows days post-anthesis (DPA) to reach the mature-green stage. The DPA to reach final fruit coloration is indicated in parentheses next to each allele. The fruit ripening stages are from USDA (https://www.ams.usda.gov/sites/default/files/media/Tomato_Visual_Aids%5B1%5D.pdf handbook).

### *RIN*-edited plants showed delayed fruit ripening

The *rin^CR1-4^* alleles showed no perceptible differences in their vegetative phenotype and onset of flowering from wild-type parent-Arka Vikas (AV). However, in fruits of all alleles, the attainment of the mature-green (MG) stage was delayed by ca. 10-13 days (**Figure 1C**). Post-MG stage, in *rin^CR1-4^*, the transition to the ripe stage was also slow. Unlike the *rin* mutant, the *rin^CR1-4^* alleles were not compromised in carotenoid accumulation. However, the ripe *rin^CR1-4^* fruits did not reach the typical red ripe (RR) stage like AV but attained the pink stage (*rin^CR4^*) and light-red stage (*rin^CR1-3^*). However, in the text and figures, the RR stage is mentioned for *rin* and *rin^CR1-4^* for brevity.

Unlike *rin*, the on-vine ripening fruits of *rin^CR1-4^*alleles showed fruit senescence, marked by skin wrinkling (**Figure S3A**). Nonetheless, the duration between the attainment of the ripe stage and the onset of fruit senescence was considerably longer in *rin^CR1-4^* alleles (**Figure S3A, B**). Except for *rin^CR4^*, the fruit firmness of the other three *rin^CR^* alleles was higher than AV but lower than *rin* (**Figure S3C**). The post-harvest shelf life of *rin^CR1-4^* fruits harvested at the RR stage was longer than the AV (AV 10 days, *rin^CR1-3^*30-35 days, *rin^CR4^* 20 days) (**Figure S4**). Unlike *rin*, the *rinCR^1-4^* alleles accumulated phytoene, phytofluene, and lycopene during ripening. Consistent with the weaker red coloration of fruits, the levels of lycopene and its precursors’ phytoene and phytofluene in on-vine ripened fruits were much lower in the *rin^CR1-4^*(**Figure 2A, Dataset S2**). Among the carotenoids downstream of lycopene, the level of lutein in *rin* was higher at the RR stage. Overall, at the RR stage, while total carotenoids in *rin* mutant were about 10% of AV, in *rinCR^1-4^* alleles, it ranged from 28-54% (**Figure S5**). Notably, the profile of carotenoid distribution in *rinCR^1-4^* alleles was somewhat intermediate between *rin* and the wild-type (**Figure S6**).

**Figure 2.**
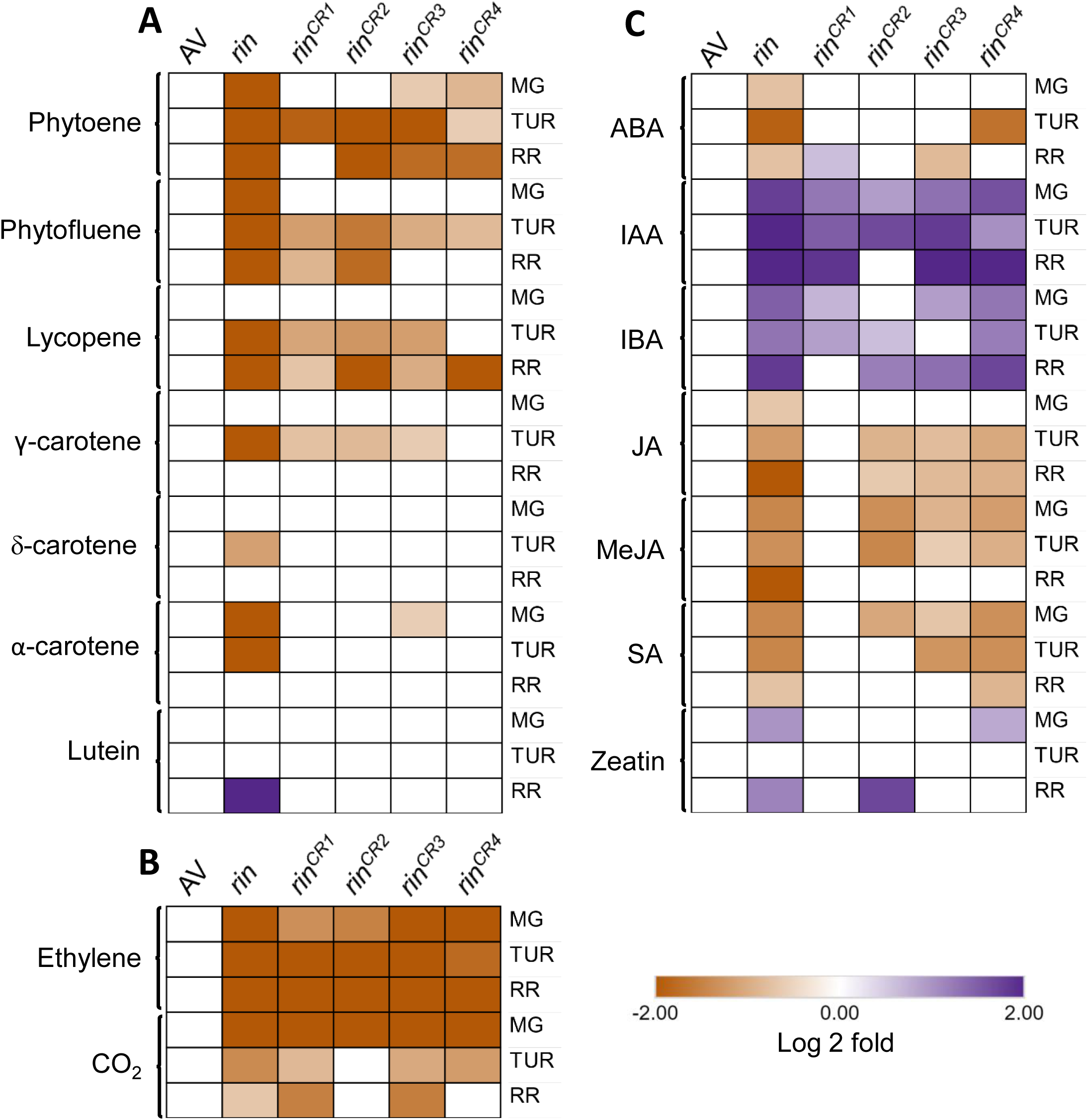
The shifts in levels of carotenoids (**A**), ethylene/CO_2_ (**B**), and hormones (**C**) *rin* and *rin^CR1-4^* fruits during ripening compared to Arka Vikas (AV). The relative changes in the carotenoids, ethylene/CO_2,_ and hormone levels at MG, TUR, and RR stages were determined by calculating the log2 fold change at respective ripening phases with reference to AV. Only significantly changed carotenoids and hormones are depicted (Log2 fold ≥ ± 0.584, p-value ≤ 0.05, n ≥ 3)). See the following datasets for detailed carotenoids (**Dataset S2**), CO_2_ (**Dataset S3**), and hormones, including ethylene (**Dataset S4**) data. *Abbreviation*: ABA-Abscisic acid; IAA-Indole acetic acid; IBA – Indole butyric acid; JA-Jasmonic acid; MeJA-Methyl Jasmonate; SA-Salicylic acid.

### Hormonal and metabolite profiles are altered in *rin^CR1-4^* fruits

In consonance with slower on-vine ripening, the ethylene and CO^2^ emission from fruits of *rin* and *rin^CR1-4^* was considerably subdued (**Figure 2B, Dataset S3-S4)**). The levels of other hormones were also analogously affected in *rin,* and *rin^CR1-4^*, particularly IAA and IBA levels, were considerably elevated. In contrast, salicylic acid (SA), methyl jasmonate (MeJA), and jasmonic acid (JA) levels were lower than AV at one or more stages, barring a few exceptions (**Figure 2C, Dataset S4**). The level of zeatin was higher in the *rin* mutant, as previously determined by bioassays (**Davey and Van Staden, 1978**), but a higher zeatin level was infrequently seen in *rin^CR^* alleles.

Like hormonal changes, the *rin^CR1-4^* alleles showed alteration of metabolic profiles, which differed from AV but were akin to the *rin*. The principal component analysis (PCA) revealed that in each *rin^CR^*allele, the PCA profile followed a trajectory different from AV but was relatively close to *rin* (**Figure S7**). Consistent with this, most amino acids and organic acid levels in *rin^CR1-4^* alleles, particularly at TUR and RR stages, were more similar to *rin* than AV (**Figure S8, Dataset S5**). The *rin* mutant and *rin^CR1-4^* alleles also showed higher °Brix and pH than AV (**Figure S9**).

### The expression of most ripening-related genes is subdued in *rin^CR^* alleles

In tomato *rin* mutant, the expression of ripening-related genes, including *RIN*, *CNR,* and those of carotenoids biosynthesis (*DXS*, *PSY1*), ethylene emission (*ACS2*, *ACS4*, *ACO1*, *ACO3*), and cell wall softening (*PG2a*, *PE1*, *EXP*, *LOXB*, *CEL*, *FSR*, *PL1*, *β*-*MAN*, *TBG4*, *XYL*) are highly reduced (**Figure 3, Dataset S6**). Consistent with this, barring a few exceptions, most genes related to the above responses were downregulated in *rin^CR1-4^* alleles. Nonetheless, the reduction in expression for most genes in *rin^CR^*alleles was less severe than *rin*. Unexpectedly, in *rin^CR2^*, the transcript levels of *RIN* were higher than AV. Among *rin^CR^* alleles, the influence of *rin^CR2^*was milder for many genes. In consonance with reduced system-II ethylene emission in *rin^CR^* alleles, while the expression of ethylene biosynthesis genes *ACS4* and *ACO1* was low, the expression of *ACO3* and *ACS2* was upregulated at the RR stage. A major deviation was higher expression of *FSR*, *PG2a,* and *PE1* in *rin^CR1-4^* alleles compared to reduced expression in *rin*. Overall, the influence of *rin^CR1-4^* alleles on gene expression though largely reminiscent of *rin* but was not exactly analogous.

**Figure 3.**
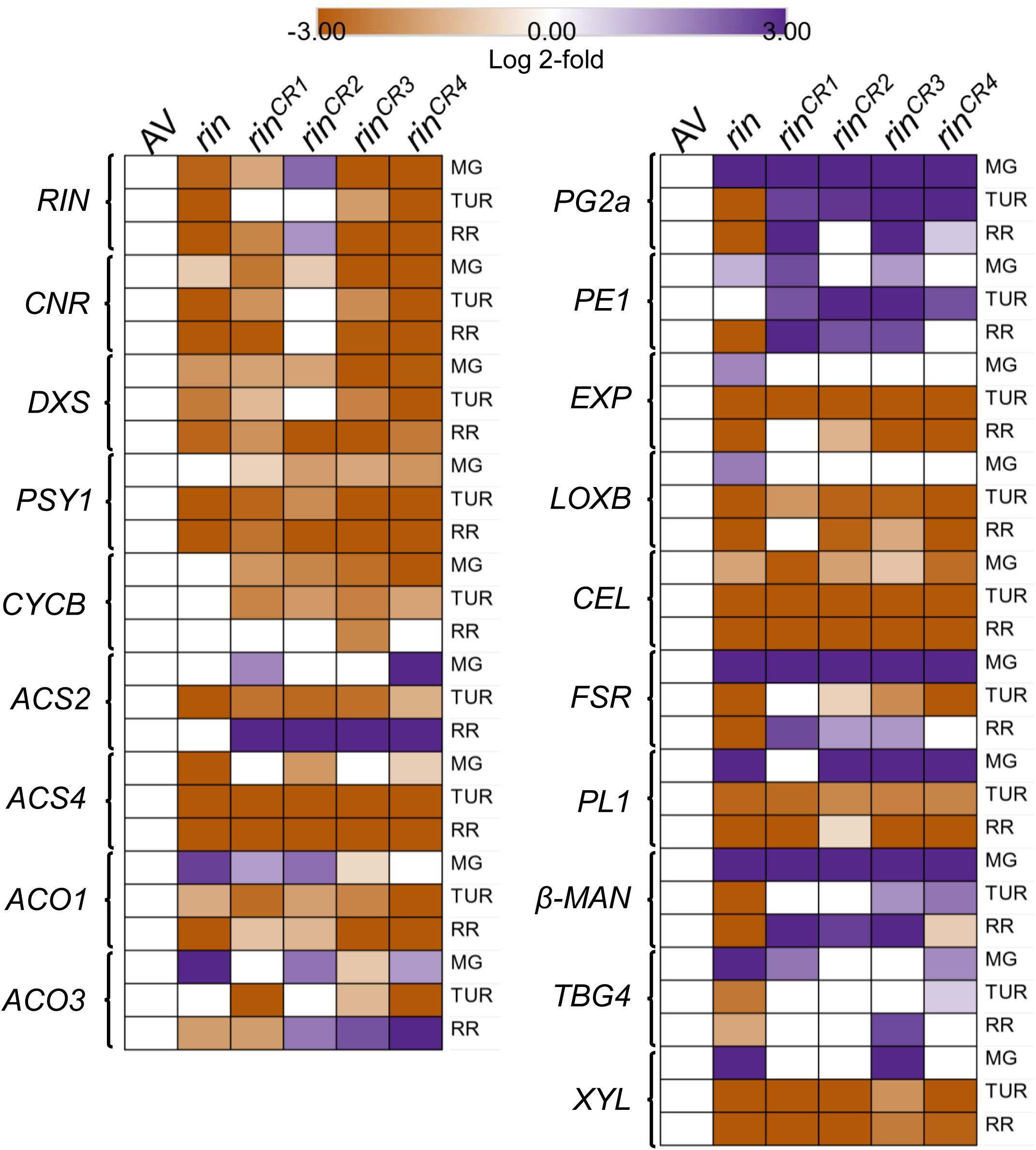
The expression of ripening-related (**A**) and cell-wall-related (**B**) genes in *rin* and *rin^CR1-4^* (T_2_) fruits during ripening compared to Arka Vikas (AV). The relative changes in gene expression at MG, TUR, and RR stages were determined by calculating the log2 fold change at respective ripening phases with reference to AV. Only significantly changed transcripts are depicted (Log2 fold ≥ ± 0.584, p-value ≤ 0.05, n ≥ 3)). See **Dataset S6** for detailed gene expression data. Gene abbreviation: *ACS2* -1-amino-cyclopropane-1-carboxylate synthase 2, *ACS4* -1-amino-cyclopropane-1-carboxylate synthase4, *ACO1* -1-amino-cyclopropane-1-carboxylate oxidase1, *ACO3* -1-amino-cyclopropane-1-carboxylate oxidase3, *RIN* -Ripening inhibitor, *CNR* -colorless non-ripening, *DXS* -1-deoxy-D-xylulose 5-phosphate synthase, *PSY1* -Phytoene synthase1, *CYCB* - Lycopene β-cyclase, *CEL* -Cellulase, *EXP1* -Expansin 1, *PG2a* -Polygalacturonase 2A, *PE1* -Pectin esterase 1, *FSR* -Fruit shelf-life regulator, *PL* -pectate lyase, *β-MAN* -β-mannosidase, *TBG4* - tomato β-galactosidase 4, *XYL* -β-D-xylosidase, *LOXB* -Lipoxygenase B.

### Introgression of *Nps1* augments carotenoids in *rin^CR^* alleles

The *rin* mutation has been used in the heterozygous state to increase the post-harvest shelf life of tomato fruits compared to normal-ripening cultivars (**Buescher et al., 1976**). However, the gain of higher shelf life in the above hybrids is marred by the loss of fruit flavour and reduced lycopene content (**Kopeliovitch et al., 1982**). We sought to overcome this drawback by crossing *rin^CR^* (*r/r*) alleles with *Nps1* (*N/N*), a dominant *phototropin1* mutant, which upregulates the carotenoids in red-ripe fruits (**Kilambi et al., 2021**). We crossed *Nps1* with two diverse *rin^CR^* alleles, viz *rin^CR1^*(in-frame single amino acid deletion) and *rin^CR3^* (protein truncation) (**Figure S10-11, Table S3**). Consistent with the dominant nature of *Nps1*, *Nps1/rin^CR1^,* and *Nps1/rin^CR3^* hybrids, both in homozygous and heterozygous conditions, had 4.4-7.1-fold higher total carotenoids in RR fruits than respective *rin^CR^* alleles (**Figure S12A-B, Dataset S7**).

In the four double hybrids, five health-promoting nutraceuticals, phytoene, phytofluene, lycopene, β-carotene, and lutein, are significantly upregulated (**Figure 4, Dataset S7**). The increase in carotenoids depended solely on *Nps1* introgression, as *rin^CR^* heterozygotes had carotenoid levels like WT homozygote and AV parental line, and *rin^CR^* homozygotes had levels like *rin^CR^*parental lines. The increase in carotenoids in *Nps1/rin^CR^* double hybrids was not correlated with the expression of carotenogenesis genes such as *CNR*, *DXS*, *PSY1,* and *CYCB*, a feature reported in the *Nps1* mutant (**Figure S13, Dataset S8**) (**Kilambi et al., 2021**). Since *Nps1/rin^CR^* double hybrids with *Nps1* and respective *rinCR* alleles had higher carotenoids in homozygous or heterozygous states, we confined further investigations to these four hybrids (*N/N*:*r/r*, *N/n*:*R/r*, *N/N*:*R/r*, *N/n*:*r/r*) in the F_2_ generation.

**Figure 4:**
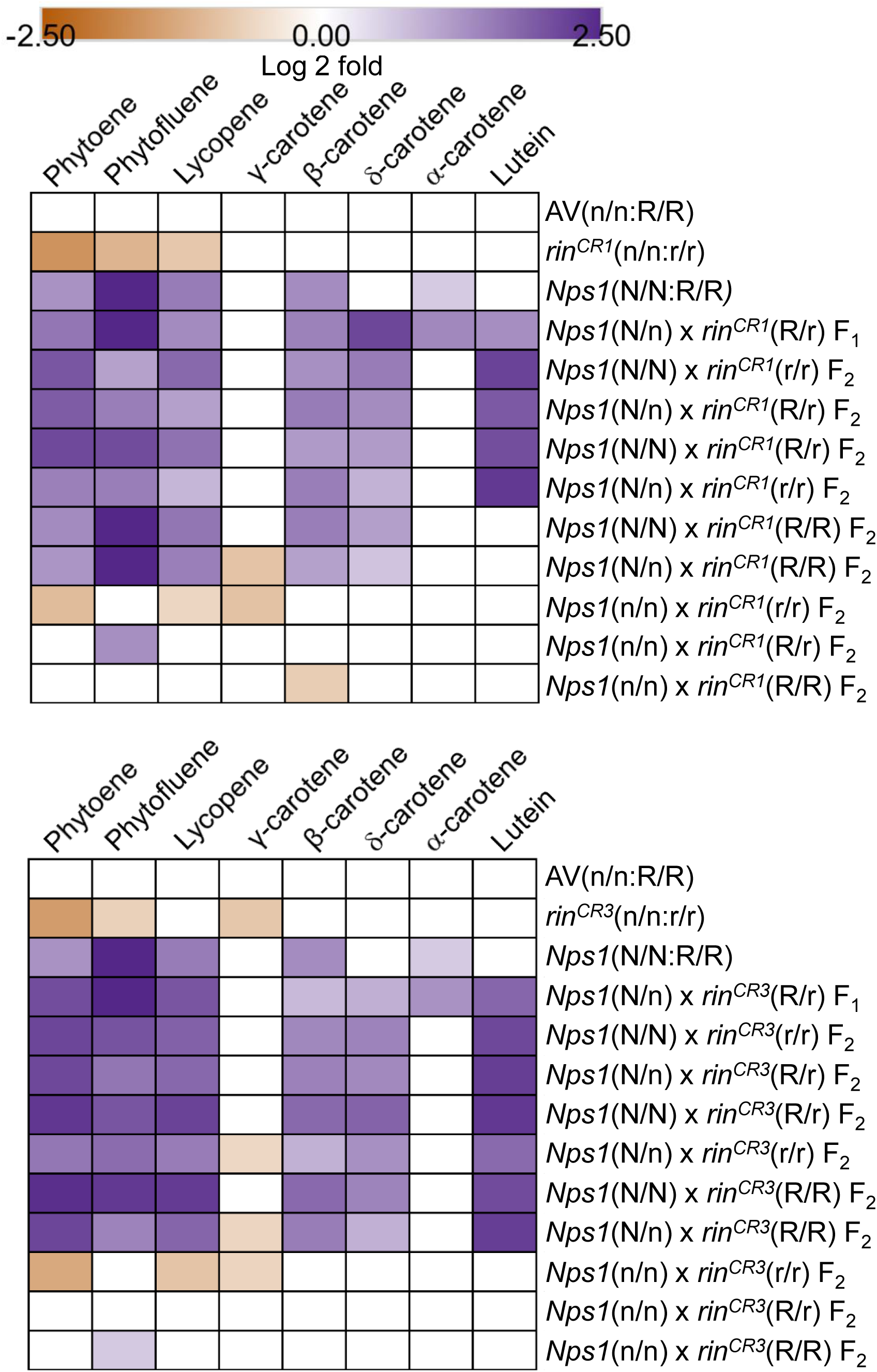
The levels of different carotenoids in *rin^CR1^* (**A**) and *rin^CR3^* (**B**) mutants (*n/n:r/r*) and their genetic segregation after crossing with the *Nps1* (*N/N:R/R*) mutant. The carotenoids were analysed at the RR stage. Note that *Nps1* (*N/N*), being a dominant mutation, enhances carotenoids in homozygous (*N/N*) and also heterozygous (*N/n*) states in *rin^CR1^*and *rin^CR^* alleles and in wild-type (R/R). The homozygous F_2_ lines bearing the wild-type allele (*n/n:R/R*) have carotenoids like Arka Vikas (AV), and the one bearing the *rin^CR1^* or *rin^CR3^* (*n/n:r/r*) is like their respective allele. Only significantly changed carotenoids with reference to Arka Vikas (AV) are depicted (Log2 fold ≥ ± 0.584, p-value ≤ 0.05, n ≥ 3). See **Dataset S7** for individual carotenoid levels and significance.

### *Nps1*/*rin^CR^* hybrids have extended off-vine shelf life

We next examined whether, in addition to the upregulation of carotenoids, the *Nps1/rin^CR^*F_2_ hybrids also displayed extended shelf life. The onset of the MG stage was delayed in respective double hybrids (36 DPA AV; 47-48 DPA *Nps1/rin^CR1^*; 48-52 DPA *Nps1/rin^CR3^*), similar to *rin^CR^* parental lines (**Table S4, S5**). Furthermore, post-MG stage, the time taken to reach the RR stage was longer in homozygous (r/r) than in heterozygous (R/r) lines (7 days AV; 7 days (R/r), 12 days (r/r) *rin^CR1^*; 12 days (R/r), 16 days (r/r) *rin^CR3^*). Like the parental *rin^CR^* lines, the onset of fruit senescence in *Nps1*/*rin^CR^* hybrids was considerably delayed (10 days AV, 50-65 days *Nps1*/*rin^CR1^*, 55-75 days *Nps1*/*rin^CR3^*) (**Figure S14-15**). Consistent with delayed on-vine senescence of the *Nps1/rin^CR^* F_2_ hybrids, the expression of the majority of genes related to cell wall softening (*PE1*, *EXP*, *LOXB*, *CEL*, *FSR*, *PL1*, *β*-*MAN*) were downregulated, barring *PG2a*, *TBG4*, and *XYL* (**Figure S13, Dataset S8**). Unlike cell wall softening genes, the expression of genes related to ethylene biosynthesis did not correlate to delayed on-vine fruit senescence (**Figure S13**). The post-harvest shelf life of *Nps1*/*rin^CR1^* and *Nps1/rin^CR3^* fruits harvested at the RR stage was longer in F_2_ hybrids than the AV [AV 10 days, *Nps1/rin^CR1^* (*N/N:r/r*) 15 days, *Nps1/rin^CR3^* (*N/N:r/r*) 20 days] (**Figure S16A-B**).

### Hormonal profiles of F_2_ hybrids show the dominance of both *Nps1* and *rin^CR^* alleles

The RR fruit of *rin^CR3^* displayed lower ABA and increased levels of IAA and IBA, a feature partially shared by *rin^CR1^* with higher IAA. Remarkably, the F_2_ hybrids bearing the *rin^CR3^*allele, too, showed lower ABA and increased IAA and IBA levels (**Figure 5**, **Dataset S9**). The F_2_ hybrids of *rin^CR1^* also showed a similar influence on ABA, IAA, and IBA levels, albeit with few gaps. Notably, the downregulation of IAA observed in *Nps1* was dominantly inverted to upregulation in *rin^CR1^* and *rin^CR3^* hybrids. A similar dominance on hormonal levels was observed for SA and zeatin modulated by *Nps1*. Barring *rin^CR^* F_2_ hybrid with *N/n:r/r* genotype, the reduced level of SA and zeatin was observed in other combinations.

**Figure 5:**
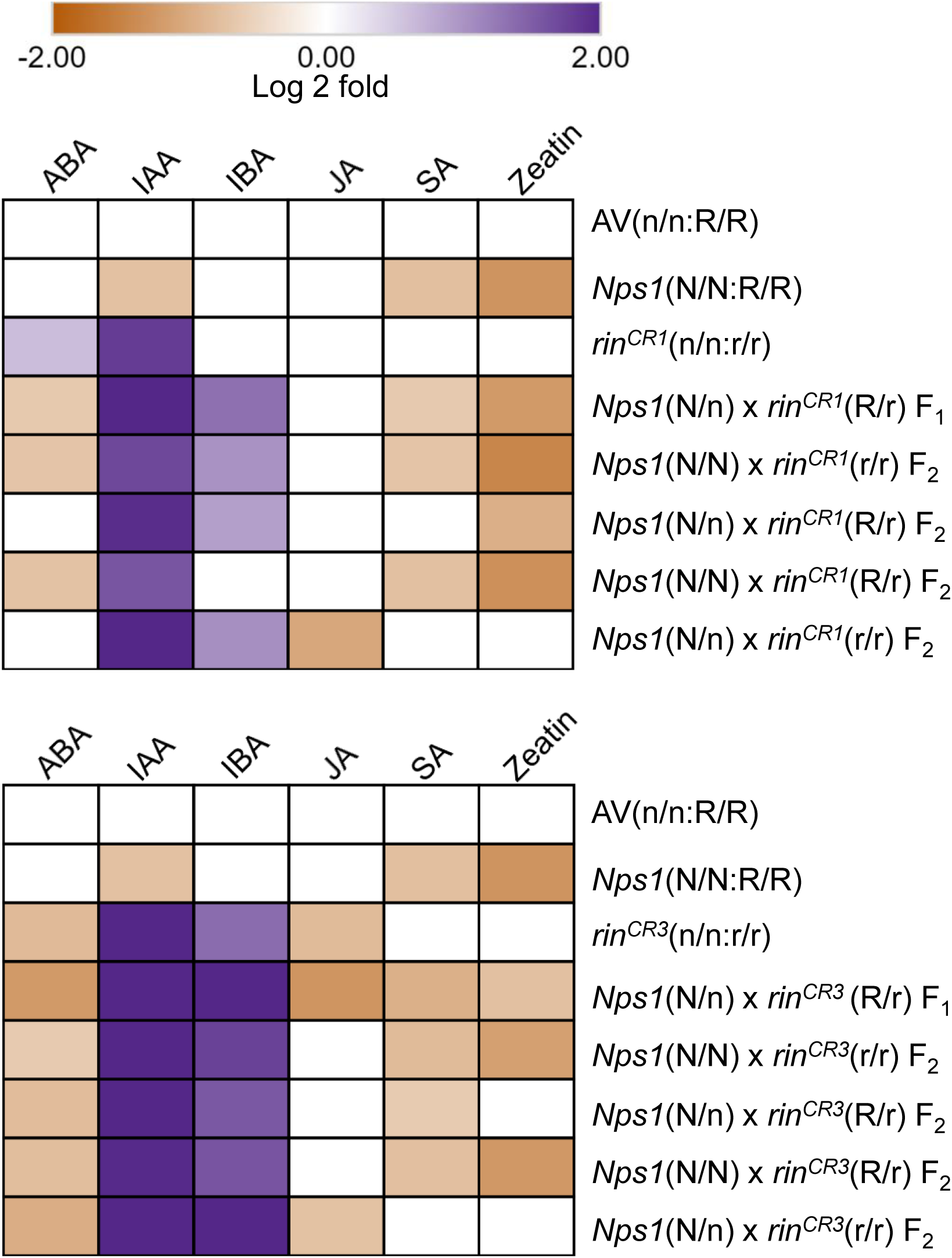
The levels of different hormones in *rin^CR1^* (**A**) and *rin^CR3^* (**B**) mutants (*n/n:r/r*) and their genetic segregation after crossing with the *Nps1* (*N/N:R/R*) mutant. The hormones were analysed at the RR stage. The dominant influence of the *rin^CR3^* allele on ABA, IAA, and IBA persists in homozygous (*r/r*) and heterozygous conditions (*R/r*), even in the presence of the *Nps1* allele. Similar to the *rin^CR3^* allele, the *Nps1* dominantly influences SA and zeatin levels in F_2_ plants, barring *Nps1*(*N/n*) x *rin^CR3^*(*r/r*) combination. Only significantly changed metabolites with reference to Arka Vikas (AV) are depicted (Log2 fold ≥ ± 0.584, p-value ≤ 0.05, n ≥ 3). See **Dataset S9** for individual hormone levels and significance. *Abbreviation*: ABA-Abscisic acid; IAA-Indole acetic acid; IBA - Indole butyric acid; JA-Jasmonic acid; SA-Salicylic acid; AV-Arka Vikas.

### *Nps1*/*rin^CR^* hybrids show intermediate metabolite profiles

Given that *Nps1*/*rin^CR^* hybrids displayed significant upregulation of carotenoids in ripe fruits and shift in hormonal profiles, we next compared whether the primary metabolome of these hybrids was also altered. The principal component analysis (PCA) of the metabolome of RR fruits revealed very distinct patterns. For *Nps1* X /*rin^CR1^* crosses, the PC of parents *Nps1*, AV, and *rin^CR1^* was distinctly separated. Interestingly, the PC of all four combinations of *Nps1*/*rin^CR1^* hybrids were in close proximity of each other and closer to AV. Likewise, for *Nps1* X *rin^CR3^* crosses, a more compact closeness was observed for PCs of *Nps1/rin^CR3^* hybrids (**Figure S17**).

Consistent with the proximity of PCs of *Nps1*/*rin^CR1^*and *Nps1*/*rin^CR3^* hybrids, the individual metabolite profiles of both hybrids closely matched in all four F_2_ double hybrids. Of 55 significantly affected metabolites in hybrids, 23 were uniquely upregulated (8) or downregulated (15) only in F_2_ double hybrids, not in *rin^CR1^* or *rin^CR3^* alleles or even in the *Nps1* parent. Of the remaining metabolites, 20 shared a pattern (9 Up, 11 down) with *rin^CR1^* and *rin^CR3^*, and *Nps1* parent, while 13 metabolites had a pattern of metabolite change opposite to that of *rin^CR^* alleles and *Nps1* parent (**Figure 6, Dataset S10**).

**Figure 6:**
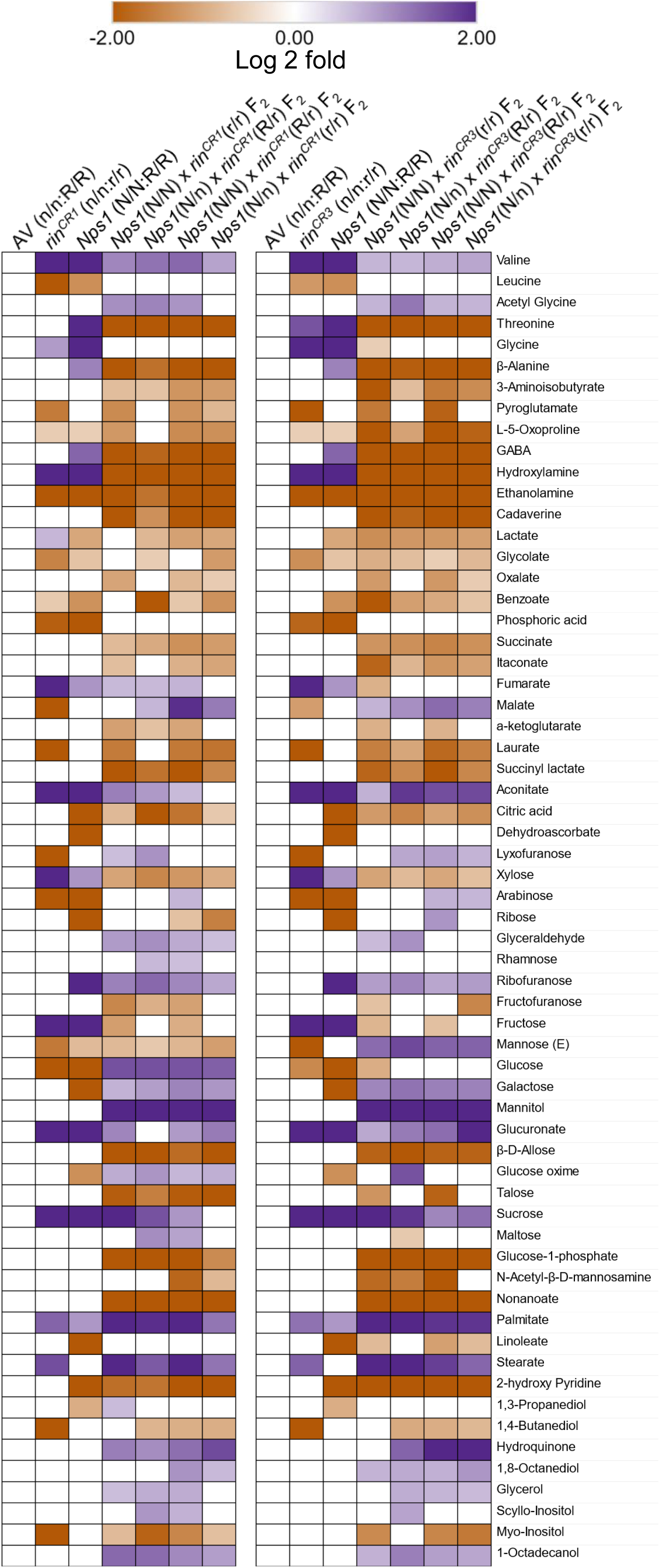
The levels of metabolites in *rin^CR1^* (left) and *rin^CR3^* (right) mutants (*n/n:r/r*) and their genetic segregation after crossing with the *Nps1* (*N/N:R/R*) mutant. The metabolites were analyzed at the RR stage. The individual metabolite profiles of *rin^CR1^* and *rin^CR3^* closely matched the profiles of all four F_2_ double hybrids. Only significantly changed metabolites with reference to Arka Vikas (AV) are depicted (Log2 fold ≥ ± 0.584, p-value ≤ 0.05, n ≥ 5). See **Dataset S10** for individual metabolite levels and significance. AV-Arka Vikas.

Consistent with high °Brix values observed in *rin^CR^*alleles, the sucrose levels were higher in *rin^CR^* parents and dihybrid F_2_ plants. However, on average, the levels of the majority of metabolites were lower in dihybrid F_2_ plants than in the AV. In particular, the metabolites related to the TCA cycle, such as citrate, α-ketoglutarate, succinate, and itaconate, were downregulated, except for malate, which was upregulated. Similarly, the amino acids and their derivatives derived from the TCA cycle and glycolysis, such as threonine, oxoproline, GABA, and hydroxyproline, also showed reduced accumulation in all four dihybrids F_2_ fruits. The upregulation of sugar alcohols such as mannitol may be related to the extended shelf life of *Nps1/rin^CR^* hybrids.

### *Nps1* homozygous *rin^CR^* hybrids boost flavour-related volatiles

Hybrids between the *rin* mutant and normal-ripening lines have been utilized commercially to produce tomatoes with longer shelf life. However, the red ripe fruits of these hybrids with the *rin* gene have reduced levels of carotenoids and volatiles associated with flavour and aroma (**Baldwin et al., 1995; Kitagawa et al., 2005**). Therefore, consumer and tasters panels ranked the *rin* hybrids as inferior in quality to normal tomatoes (**Baldwin et al., 1995**). Contrasting to the above *rin* hybrids, the *Nps1/rin^CR^* hybrids showed significant upregulation of flavour-related volatiles. Intriguingly, the above upregulation was confined to the genic combination (*N*/*N:r*/*r*; *N*/*N:R*/*r*) bearing homozygous *Nps1* (*N*/*N*). Volatiles originating from fatty acids, such as 1-penten-3-ol, pentanal, and pentanol, which provide fruity notes to tomatoes, and are appreciated by consumers, were upregulated (**Shen et al., 2014**). Conversely, the compounds linked with undesirable flavours, such as 2-hexanol; 2,4,-hexadienal, are downregulated. The carotenoid-derived volatile 6-methyl-5-hepten-one, responsible for the tomato-like flavour and sweet floral aroma, was upregulated in the *Nps1* homozygote (**Figure 7, Dataset S11**).

**Figure 7:**
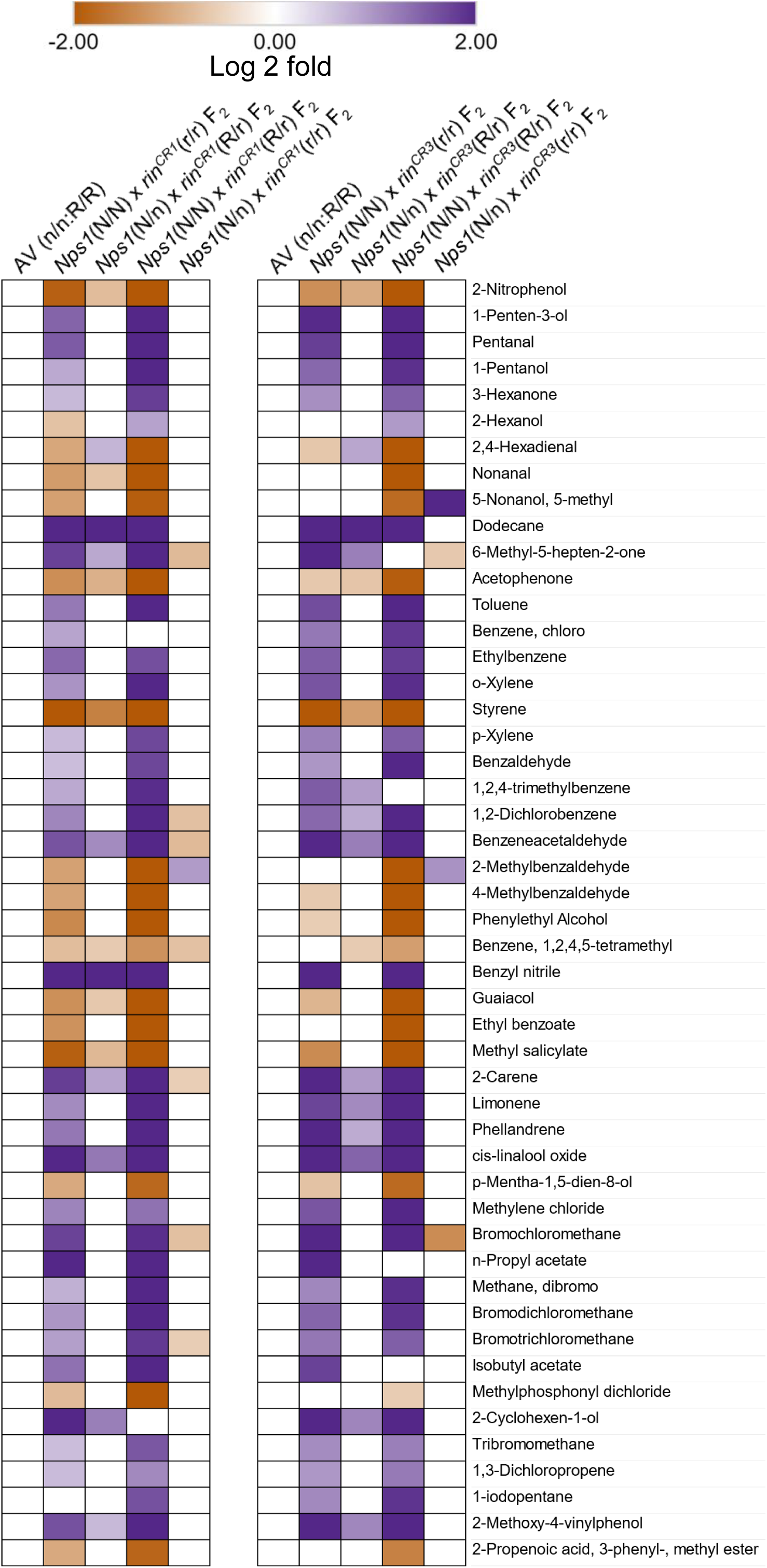
The levels of volatiles in RR fruits of four dihybrids of *Nps1* X *rin^CR1^* (**Left**) and *Nps1* X *rin^CR3^* (**right**) crosses. The upregulation of volatiles by *Nps1* is prominently seen only in homozygous (*N/N*) conditions. Only significantly changed volatiles with reference to Arka Vikas (AV) are depicted (Log2 fold ≥ ± 0.584, p-value ≤ 0.05, n = 5)). See **Dataset S11** for individual volatile levels and significance.

Remarkably, the *Nps1* homozygote reversed the downregulation of 6-methyl-5-hepten-one observed in the *rin* homozygote (*r*/*r*), substantiating the dominance of the *Nps1* homozygote on the flavour profiles. The *Nps1* homozygotes upregulated benzaldehyde associated with a sweet smell. Significantly, the methyl salicylate and guaiacol associated with the negative taste of tomato were downregulated in *Nps1* homozygotes (**Tieman et al., 2017**). Among the terpenoids, the limonene and linalool oxide levels with a lemon/orange flavour were upregulated in *Nps1* homozygotes. To sum up, the introgression of *Nps1* significantly reversed the negative effect of *rin^CR^*alleles on flavour/aroma profiles.

## Discussion

Fruit ripening in tomato is influenced by a myriad of factors, including genetics, hormones, and environmental conditions. At the pinnacle of these factors, it was believed that the RIN acts as a master regulator of ripening initiation. The above role of RIN in tomato fruit ripening is now revised. The analysis of gene-edited *RIN* alleles revealed that the primary function of RIN is to suppress fruit overripening. Nonetheless, the gene-edited alleles of *RIN* retain several negative traits associated with *rin,* such as reduced accumulation of carotenoids and decline in aroma/flavour-related volatiles. Using newer alleles of *RIN*, we show that the introgression of a dominant phototropin mutation, *Nps1*, can mitigate the negative traits.

### RIN K domain gene-edited alleles show distinct phenotypes

The K domain in MADS-box proteins is responsible for binding to the promoters of target genes and protein-protein interactions (**Lai et al., 2019**). Differing from *rinG3* (3-10%; BR+7 days) (RIN null mutants, **Ito et al., 2017**), *rinG2* (12%, BR+7) (Intact K Domain **Ito et al., 2020**) and *RIN-CRISPR* (7-10%; BR+10 days) (truncated RIN protein 1-∼78 AA **Li et al., 2020**) gene-edited alleles (**Fig SX**), the *rin^CR^* alleles showed accumulation of ca. 28-54% lycopene of AV (100%) at RR. However, the above differences in the lycopene levels between our and other studies may ensue as we measured carotenoids at the RR stage (AV 10 days, *rin^CR1-3^*30-35 days, *rin^CR4^* 20 days from MG). **Li et al. (2020)** observed that RIN-CRISPR fruits became orange but not red over a longer ripening duration. Considering that the *rin^CR^* alleles have extended shelf life but reduced lycopene levels, these two processes seemingly are unlinked. Similarly, **Ito et al. (2020)** observed that in *rinG2* alleles, lycopene accumulation and softening were independently regulated.

The reduced expression of carotenogenesis genes *DXS*, *PSY1,* and *CYCB* in *rin^CR^* alleles is in conformity with subdued carotenoid synthesis. Interestingly the distribution profile of β-carotene, antheraxanthin, ẟ-carotene, and α-carotene in *rin^CR^*alleles is more akin to *rin*. Remarkably, the *rin^CR^* alleles promoted carotenoid synthesis with a somewhat intermediate carotenoid profile between wild-type and *rin*. Considering that *rin^CR^* alleles show a carotenoid profile more similar to *rin*, a functional *RIN* is necessary to execute a ripening-specific carotenoid profile.

The partial semblance of action of *rin^CR^* alleles to *rin* is also observed in the ripening induction. However, unlike *rin*, the *rinG3, rinG2*, and *RIN-CRISPR* alleles show no or only a minor delay in the onset of the breaker stage **(Ito et al., 2020; Li et al., 2020).** In contrast, the onset of the breaker stage is delayed by >10 days in *rin^CR^* alleles, suggesting that conversely to carotenoid synthesis, the partial/altered K domain negatively affects the ripening initiation and progression. The near doubling of time in *rin^CR^* alleles to attain the RR stage is consistent with this view. In conformity with the above view, an EMS *rin-2* mutant in the K domain (Leu^112^→Pro^112^) too displays a 14-day delay in ripening initiation and increased duration (9-days) to complete ripening (**Hoogstrate et al., 2014)**. Since another EMS *rin-3* mutant at Q^88^, with intact MADS domain but almost no K domain, shows only a 2-day delay in ripening initiation (**Hoogstrate et al., 2014)**, similar to that reported for *RIN-CRISPR* (**Li et al., 2020**) reaffirms the purported role of K domain in determining the onset of ripening.

In MADS-box proteins, the K domain plays a prominent role in forming DNA-binding complexes with other MADS proteins. The RIN protein reportedly forms complexes with TAGL1, FUL1, and FUL2. The truncated RIN protein encoded by the *rinG2* allele, which has an intact K domain but truncated C domain, forms tetramers or heteromers with FUL1, FUL2, or TAGL1, which bind to promoters but lack transcription activation (**Ito et al., 2020**). Like *rinG2* alleles, *rin^CR^* alleles, too, show extended shelf life, albeit the duration is shorter than *rinG2*. Considering the above, it is plausible that the partial K domain in *rin^CR1-2^* and the altered K domain in *rin^CR3-4^*may form complexes with other MADS-proteins and influence the expression of the target genes by acting as a repressor. In turn, it may have caused subdued carotenoid synthesis and delayed onset of ripening in *rin^CR^* alleles.

### Metabolic and hormonal homeostasis of *rin^CR^* alleles is more akin to *rin*

Metabolically tomato ripening is an intense process triggered by system II ethylene synthesis accompanied by increased CO_2_ emission. Consistent with delayed ripening, both ethylene and CO_2_ emission from *rin^CR^* alleles are subdued and more similar to *rin*. Intriguingly, while the *ACS* and *ACO* expression in *rin^CR^* alleles at TUR is suppressed like *rinG1-3* (**Ito et al., 2017, 2020**) and *RIN-CRISPR* alleles (**Li et al., 2020**), at RR stage *ACS2* and *ACO3* show higher expression than AV. Hitherto the RR stage (BR+18-22 days in *rin^CR^* alleles) signifying ripening completion was not examined in other *RIN* gene-edited alleles (**Ito et al., 2017** BR+4**; Ito et al., 2020** BR+7**; Li et al., 2020** BR+10). The lower levels of lycopene detected in *rinG1-3* and *RIN-CRISPR* than in *rin^CR^* alleles may be due to an incomplete ripening stage used for these alleles.

The similarity of *rin* and *rin^CR^* alleles was not limited to ethylene and manifested in other phytohormones. The similarity in IBA and IAA levels at all stages of ripening to *rin* reinforces the view that *rin^CR^*alleles are more similar to *rin* than AV. The levels of other hormones in *rin^CR^* alleles were, in general, more alike to *rin* than AV. The parallelism of hormonal profiles between *rin* and *rin^CR^* alleles also manifests in metabolic profiles, particularly at the TUR and RR stages. The above parallelism implies that though *rin* represses ripening as a fusion protein with macrocalyx, the gene-edited *rin^CR^* alleles function more like weaker *rin* alleles.

Consistent with this view, the fruits of *rin^CR^* alleles show typical features of *rin* mutant, such as the delayed onset of BR, a longer time to reach RR, and prolonged delay in the onset of senescence in on-vine ripened fruits. In a like manner, the post-harvest shelf life of fruits of *rin^CR^* alleles is also much longer than AV. In conformity, with delayed on-vine and post-harvest fruit spoilage, the expression of most cell wall softening-related genes was subdued in *rin^CR^*alleles like *rin* and other *RIN*-edited alleles (**Ito et al., 2017; Li et al., 2020**). Lowered expression of cell wall softening-related genes may contribute to delayed senescence of *rin^CR^* alleles. Alike our study, a high expression of *PG2a* was also observed in *rinG3, rinG1* (**Ito et al., 2017**), and *RIN-CRISPR* (**Li et al., 2020**) edited lines at a TUR-equivalent stage, indicating loss of RIN function upregulates *PG2a*.

### ***Nps1* introgression boosts carotenoids in *rin^CR^* fruits**

Though the introgression of *rin* in diverse cultivars augments the shelf life, the hybrid fruits are bereft of the carotenoids and aroma (**Kopeliovitch et al., 1982; Kitagawa et al., 2005).** Since F_1_ hybrids of *rinG3* alleles also have lower carotenoid levels than the parent (**Ito et al. 2017**), it can be surmised that the WT *RIN* allele cannot fully restore the carotenoids biosynthesis in *rinG3* hybrids. In contrast to *rin*, which accumulates only traces of carotenoid precursors and lycopene (**Gupta et al., 2022; Dataset S2)**, the *rin^CR1-4^* alleles accumulated higher levels of phytoene, phytofluene, and lycopene than *rin*, albeit at a much lower level than AV. In *rin,* the accumulation of β-carotene, but not lycopene, is ascribed to higher expression of the lycopene β-cyclase (*CYCB*) gene (**Ito et al. 2020)**. However, the attempt to elevate the carotenoid levels in *rin* by generating a loss-of-function mutation in the *CrtL1* (lycopene β-cyclase-*CYCB*) gene in *rinG2* fruits in the heterozygous state (homozygous were lethal), only moderately boosted the lycopene levels (12%◊37% lycopene) (**Ito et al. 2020)**.

In contrast to **Ito et al. (2020)**, the homozygous introgression of *Nps1* into *rin^CR^* alleles significantly increased lycopene levels compared to the AV (*rin^CR1^*: 330%, *N/N:r/r; rin^CR3^*: 350%, *N/N:r/r*). Even in the heterozygous *Nps1* hybrids, though the level of lycopene was lower than homozygous *Nps1*, it was still higher than in the wild type (*rin^CR1^*: 179%, *N/n:r/r*; *rin^CR3^*: 283%, *N/n:r/r*). The above difference in upregulation entails that *Nps1* affects the carotenoid level in a semidominant fashion. Since in homozygous F_2_ *rin^CR1^* (116%, *n/n:r/r*) and *rin^CR3^* (79%, *n/n:r/r*) lycopene level was akin to the respective *rin^CR^* parental line; it can be construed that the enhanced lycopene level resulted from *Nps1* introgression. Moreover, the exclusive detection of ζ-carotene, the upregulation of phytoene and phytofluene-precursors to lycopene, along with β-carotene and δ-carotene reinforce that *Nps1* introgression (*N/N* or *N/n*), boosted overall carotenoid biosynthesis in *rin^CR^* (*r/r* or *R/r*) alleles.

The accumulation of chromoplast-specific carotenoids in tomato is generally associated with the upregulation of *PSY1* and downregulation of *CYCB* transcripts (**Stanley and Yuan, 2019**). While the *Nps1/rin^CR1^* and *Nps1/rin^CR3^*homo-/hetero-zygotes showed increased carotenoid levels, it was not associated with increased expression of *PSY1* and decline in *CYCB*. Even the expression of system II ethylene biosynthesis genes, *ACO1* (barring *rin^CR1^*), *ACO3*, *ACS2,* and *ACS4,* were not affected in *Nps1/rin^CR^* homo-/hetero-zygotes. In *Nps1,* increased carotenoid levels are not related to system II ethylene biosynthesis and gene expression (**Kilambi et al., 2021**). Therefore, it is likely that the *Nps1* introgression increased carotenoids through a mechanism other than the modulation of carotenogenesis- and ethylene-related gene expression. Based on proteome profiling of *Nps1*, **Kilambi et al. (2021**) suggested that *Nps1* seemingly elevates carotenoids due to better protein protection for carotenogenesis and increased levels of carotenoid sequestration proteins (**Kilambi et al., 2021**). Therefore, it is plausible that a proteome-related process may have elevated carotenoids in *Nps1/rin^CR^*hybrids.

### The metabolome of *Nps1/rin^CR^* hybrids is divergent from parents

Unlike carotenoids, the metabolome of *Nps1/rin^CR^* hybrids distinctly differed from either of the parental lines viz, *rin^CR^* or *Nps1* and even from its progenitor line AV. This difference was prominently reflected in PCA, where profiles of four hybrids of *Nps1/rin^CR1^*and *Nps1/rin^CR3^* were clustered together, while those of *rin^CR^* alleles or *Nps1* or AV were separated from hybrids and each other. The above principal component clustering in hybrids is consistent with the view that hybridization leads to altered gene expression, enzyme activity, and metabolic pathways (**Meyer et al., 2012**). Thus, the metabolome of the hybrid is distinct from either parent, with very few metabolites in the *Nps1/rin^CR^*hybrids overlapping with either parent.

Considering that we analysed metabolome in *Nps1/rin^CR^* hybrid fruits at RR stage (∼ 60 dpa), our data can be best compared with **Osario et al. (2011),** wherein they compared the metabolome of *rin* with *nor*, *Nr,* and fully ripe parent Alisa Craig (57 dpa). While *Nr* and *nor* showed substantially lower levels, the cell-wall-related metabolites in *rin* were nearly similar to Alisa Craig. Similar to **Osario et al. (2011)**, no defined change in cell-wall-related metabolites like rhamnose, ribose, and arabinose was found, except for increased ribofuranose in *Nps1/rin^CR^* hybrids. Nevertheless, consistent with increased post-harvest shelf life and delayed on-vine senescence of *Nps1/rin^CR^* hybrids, similar to parental lines, the number of genes associated with cell wall degradation, such as *CEL*, *EXP1, PE1*, *FSR*, *PL*, and *β*-*MAN* were downregulated in *Nps1/rin^CR^* hybrids. Considering that higher expression of these genes plays a major role in cell-wall softening (**Ito et al., 2017, 2020; Li et al., 2020**), the increased shelf life of *Nps1/rin^CR^* hybrids may ensue from reduced expression of the above genes.

The ripening of tomato fruit involves the synchronization of various metabolic processes initiated by ethylene, which are also coordinated with the actions of other hormones. While the role of ethylene is well established, little is known about the function of other hormones, such as SA and zeatin, in tomato ripening (**Li et al., 2021**). Therefore, the link between the reduced levels of SA and zeatin to the ripening regulation of *Nps1/rin^CR^* hybrids remains equivocal. A feature of *Nps1/rin^CR^* hybrids was reduced levels of ABA, whose lowering is associated with reduced expression of cell wall degradation genes and the increased post-harvest life of fruits (**Sun et al., 2012; Zhang et al., 2009**). The higher auxin levels in *Nps1/rin^CR^* hybrids are in conformity with the notion that auxin functions as a repressor of climacteric fruit ripening (**Lieberman et al., 1977; Su et al., 2015**). The auxin signalling considerably declines during tomato ripening but not in the *rin* mutant **(Shin et al., 2019**). Moreover, exogenous IAA inhibits tomato ripening and downregulates cell wall-modifying genes (**Chirinos et al., 2023**). Therefore, it is plausible that the higher levels of auxin and reduced levels of ABA in *Nps1/rin^CR^* hybrids combinedly contribute to their increased shelf life.

A positive trait of *Nps1/rin^CR^* hybrids was increased sucrose levels, which may contribute to the sweetness of tomato fruits. Conversely, non-ripening mutants like *Nr*, *nor,* and *rin* have significantly lower sucrose levels than the parental line (**Osario et al., 2011).** Except for high glucose levels in *Nps1/rin^CR1^*, the levels of fructose and glucose did not differ from AV. In contrast to *rin,* which shows no significant change in TCA cycle intermediates (**Osario et al., 2011)**, *Nps1/rin^CR^* displayed a decrease in citrate, succinate, α-ketoglutarate, and itaconate levels and an increase in aconitate and malate levels. The downregulation of the above TCA cycle intermediates indicates a change in the overall metabolic flux and may, in turn, lead to respiration attenuation (**Centeno et al., 2011**). The altered levels of TCA cycle intermediates, lower levels of most amino acids, lower ABA, and higher IAA levels reflect the wide-ranging influence of *Nps1/rin^CR^* hybrids on the cellular metabolome. Seemingly, the above metabolome changes are phenotypically displayed as increased post-harvest shelf life and delayed fruit senescence in *Nps1/rin^CR^* hybrids.

### Improved volatile profiles are restricted to hybrids bearing homozygous *Nps1* (N/N)

Tomato contains a variety of volatiles that contribute to its characteristic aroma and are appreciated by consumers. The higher shelf life, along with better taste and aroma, is a desirable goal of tomato breeders. However, the introgression of *rin* by breeders to improve shelf-life in hybrids is marred by the negative impact of *rin* on tomato taste and flavour (**Kovacs et al., 2009; Osorio et al., 2020; Baldwin et al., 1995**). Even in *RIN-CRISPR* lines, the volatiles, including those associated with consumer liking, were significantly reduced (**Li et al., 2020**). In RR fruits of *rin* F_1_ plants (*rin*/+), the volatile levels were lower than in the parental lines (**Wang et al., 2020**), indicating that wild-type *RIN* is ineffective at countering the suppressive action of *rin*.

A comprehensive analysis of consumer-preferred volatiles combined with GWAS of 398 modern and heirloom varieties and *S. pimpinellifolium* revealed that the regulation of volatiles in tomato is complex and involves a network of QTLs (**Tieman et al., 2017**). The above complexity in volatile regulation is in conformity with the fact that volatiles are derived from diverse metabolites, such as amino acids, carotenoids, and fatty acids, which are influenced by multiple genes (**Li et al., 2020**). Among the regulatory QTLs, only one gene-specific locus *E8* was identified, which positively regulated methyl salicylate and guaiacol (**Tieman et al., 2017**). Given the diversity in twenty-nine volatiles that are preferred by consumers, genetically improving tomato flavour is a challenging task. Ten of these volatiles are derived from phenylalanine, leucine, and isoleucine, while another ten are contributed by linoleic and linolenic acid. In *Nps1/rin^CR^* hybrids, excepting carotenoids, the above precursor metabolites are either unaffected or downregulated.

The observed upregulation of volatiles by *Nps1*, bearing a dominant-negative mutation in *phototropin1,* is wide-ranging. This may be related to the pleiotropic effect of photoreceptors and light-signalling gene mutants on a multitude of processes during tomato ripening (**Gupta et al., 2014; Alves et al., 2020; Kilambi et al., 2021; Wang et al., 2021**). However, despite *Nps1* being a dominant-negative mutation in the phototropin1 gene, unexpectedly, its major influence on volatile levels in *Nps1/rin^CR^* hybrids was limited to plants with homozygous (*N/N*) mutation. The above feature differs from metabolites and carotenoids, where *Nps1* modulates their levels in both the homozygous (N/N) and heterozygous (N/n) states. Apparently, a single copy of *Nps1* is insufficient to elicit its response on volatiles. Although *Nps1/rin^CR^* hybrids show a massive increase in carotenoids, the upregulation of carotenoid-derived volatile 6-methyl-5-hepten-2-one is observed only in a few, with *N/n:r/r* hybrid showing a downregulation instead. In contrast, another carotenoid-derived volatile, acetophenone, is downregulated, indicating that the influence of *Nps1,* even on individual carotenoid-derived volatiles, differs and is not related to the *Nps1*-mediated increase in carotenoid levels.

An important feature was the downregulation of methyl salicylate and guaiacol in homozygous *Nps1* hybrids. These two volatiles are associated with the negative taste of tomato. Since their upregulation in tomato is attributed to the *E8* gene (**Tieman et al., 2017**), their downregulation may be related to the reduction in ethylene synthesis/signalling by *Nps1*. Additionally, other volatiles associated with undesirable flavors, such as 2-hexanol, 2,4,-hexadienal, are also downregulated. In general, the homozygous *Nps1* hybrids show an increase in several consumer-preferred volatiles associated with tomato taste/flavour, such as 6-methyl-5-hepten-2-one, 1-penten-3-ol, nonanal, benzaldehyde, along with high malate and sucrose (**Shen et al., 2014; Tieman et al., 2017**).

This entails that the regulation of volatiles in *Nps1* homozygotes is not limited to carotenoids and includes fatty acids and amino acids. Nonetheless, the restriction of volatile regulation to *Nps1* homozygotes is perplexing. It is somewhat reminiscent of haploinsufficiency, but *Nps1,* being a dominant-negative mutation, does not fit this category. It is more probable that *rin^CR^* acts as a suppressor of ripening-related metabolism, while *Nps1* functions to overcome it. Therefore, due to the antagonistic action of these two mutations on metabolism, two copies of *Nps1* are needed to exert its influence on volatiles.

### The interaction between RIN and phototropin1 signalling is not known

**Li et al. (2019)** opined that a linear model of tomato ripening control is insufficient because multiple genes regulate ripening, and instead proposed that the ripening is regulated by a spatiotemporal cross-interacting signalling network. Presently, the specific nature of the phototropin1 signalling pathway in ripening fruits and how it is affected in *Nps1* is not known. It remains to be deciphered how the dominant phototropin1 mutation overcomes the negative impacts of *rin^CR^* mutations on ripening. Unlike RIN, a nuclear-localized transcription factor, phototropin1 is a plasma and chloroplast membrane-localised protein (**Wan et al., 2008; Kong et al., 2013**); thereby, it is unlikely that the two proteins interact directly. Due to different spatial localization, it is more likely that their respective signalling pathways intersect. Gene expression profiles of *RIN* and *phototropin1* during tomato ripening show significant overlap. Consistent being a ripening enhancer, the *RIN* expression substantially increases post-MG stage, whereas *phototopin1* (Solyc11g072710) constitutively expresses right from anthesis till full ripening **(Figure S19, Shinozaki et al., 2018;** https://tea.solgenomics.net).

A likely interaction between two pathways is hinted, as the transcriptome of *rinG2* and *rin-KO* mutant revealed significantly reduced expression of *phototropin1* in *rin-KO* mutant at the P-stage compared to wild-type (**Ito et al., 2020, Supplemental Datasets**). Although the promoter region of *phototropin1* contains a potential RIN binding site (CArG box), no direct binding of RIN was found in a ChIP assay (**Fujisawa et al., 2013, Supplemental Datasets**). Taken together, it is plausible that RIN and phototropin signalling may intersect; however, the nature and extent of interaction between them remain to be explored. Notwithstanding the above, the upregulation of protein protection is suggested as a possible mechanism for the upregulation of carotenoids in *Nps1* (**Kilambi et al., 2021)**; whether a similar or different mechanism regulates volatiles remains to be investigated.

In conclusion, the introgression of *Nps1* counteracts the detrimental effects of *rin^CR^* alleles on various metabolites, carotenoids, and volatile levels. The use of *Nps1,* in combination with gene-edited *RIN* alleles, represents a promising approach for improving fruit quality with extended shelf life. Future advances in molecular cell biology may uncover the underlying mechanism of *Nps1* signalling and its interaction with *rin^CR^* alleles.

## Materials and methods

### Plant material and growth conditions

Tomato (*Solanum lycopersicum)* cultivar Arka Vikas (IIHR, Bengaluru, India) and *non-phototropic seedling 1* mutant, *Nps1* in Arka Vikas (AV) background (BC_4_F_3_) (**Kilambi et al., 2021**) were used. The tomato *rin* (LA1795) mutant was obtained from Tomato Genomics Resource Centre, Davis, USA. (https://tgrc.ucdavis.edu/Data/Acc/AccDetail.aspx?AccessionNum=LA1795). Plants were grown in the greenhouse under natural photoperiod (12-14h day, 28±1°C; 10-12h night, 14-18°C) as described by **Bodanapu et al. (2016).** The fruits of AV, *Nps1, rin*, and *rin^CR1-4^* alleles were harvested at mature-green (MG), TUR (TUR), and red-ripe (RR) stages. The attainment of uniform fruit coloration in *rin* and *rin^CR1-4^* was considered equivalent to the RR stage.

The crosses were made between *Nps1* and gene-edited *rin^CR^* (*rin^CR1^* and *rin^CR3^*) plants. The F_1_ plants were self-pollinated to generate F_2_ plants. The fruits were harvested from the first and second truss of plants. During the crossing, the zygosity of progeny for individual *rin^CR^* alleles, *Nps1*, and respective wild-type genes, at each stage was identified by PCR-RE assay (described below), Sanger sequencing (Macrogen, Korea), and CELI endonuclease assay (**Mohan et al., 2016**). In the case of *rin*, *rin^CR1-4^* alleles, and few F_2_ crosses, where fruits do not attain the RR stage, the fruits were harvested when they reached uniform fruit coloration at the light red/pink stage. The fruits were flash-frozen in liquid nitrogen and stored at -80°C until further use, as described previously (**Sharma et al., 2021**).

### Selection of target site and plasmid construction

Single guide RNA (sgRNA) of *Ripening Inhibitor* (*RIN*; Solyc05g012020) gene was selected using the CRISPR-P web tool (http://cbi.hzau.edu.cn/crispr) (**Yang et al., 2014**). To obtain better editing efficiency, sgRNA was selected based on GC content between 40-60%. The sgRNA was folded *in silico* by the RNAfold server (http://mfold.rna.albany.edu/?q=mfold/RNA-Folding-Form2.3) to check the dangling spacer and stable hairpin structure. The human codon-optimized CAS9 with 2x35S CaMV promoter, and *NOS* terminator was amplified from pAGM4723 plasmid (Addgene# 49772) using *Kpn*I forward *Pac*I reverse primers. The amplified products were cloned into the pCAMBIA2300 vector. The gRNA expression cassette was developed through overlap extension PCR. Briefly, two long primers of sgRNA expression cassette of 130 bp were annealed through overlap extension PCR. Using *Pac*I forward and *Spe*I reverse primers, the sgRNA expression cassette was amplified and cloned into the pCAMBIA2300-CAS9 vector. The final vector pCAMBIA2300-CAS9-*RIN* was sequence-verified. The primers’ information is listed in **Table S6.**

### Transformation and analysis of genome-edited lines

*Agrobacterium tumefaciens*-mediated transformation of tomato cultivar Arka Vikas (AV, https://www.iihr.res.in/tomato-arka-vikas) was performed according to Van Eck et al. (2019). Genomic DNA was isolated from plants/seedlings as described in Sreelakshmi et al. (2010). To detect gene editing, a PCR-RE assay was performed. The PCR product encompassing about 400 bp region of gRNA target site was digested with *Tai*I enzyme. The edited lines were identified by lack of restriction enzyme site in PCR-product on agarose gel electrophoresis and confirmed by cloning and Sanger sequencing. Off-targets were identified using CRISPR-P and CRISPOR web tools (http://cbi.hzau.edu.cn/crispr; http://crispor.tefor.net) (**Yang et al., 2014; Haeussler et al., 2016**) (**Table S2**). Based on the high CFD off-target score (**Haeussler et al., 2016**), four potential off-targets were analysed using Sanger sequencing.

### Profiling of phytohormones, carotenoids, and metabolites

The phytohormone levels were determined from red-ripe fruits using Orbitrap Exactive-plus LC-MS following the protocol described earlier (**Pan et al., 2010; Bodanapu et al., 2016)**. Carotenoid profiling was carried out following the procedure of **Gupta et al. (2015).** Metabolite analysis by GC-MS was carried out by a method modified from **Roessner et al. (2000)** described in **Bodanapu et al. (2016).** The protocols used for °Brix, fruit firmness, and pH were as outlined in **Gupta et al. (2014),** ethylene emission (**Kilambi et al., 2013),** and CO_2_ emission (**Sharma et al., 2021)**. Only metabolites with a ≥1.5-fold change (log2 ± 0.584) and p-value ≤0.05 relative to the wild-type control were considered to be changed.

### On-vine senescence and post-harvest shelf life

The on-vine fruit development was monitored from the day of post-anthesis through different ripening stages. For the fruit senescence study, the fruits were left on the vine and photographed. For off-vine shelf-life studies, the freshly harvested RR fruits were incubated at 24±2°C under natural day/night conditions (16 h light and 8 h dark). For both on-vine and off-vine studies, the appearance of wrinkling on fruit skin was taken as a mark of senescence initiation (**Sharma et al., 2021)**.

### Real-time PCR

The RNA was isolated from the fruit pericarp using TRI reagent (Sigma-Aldrich, St. Louis, MI, USA), and 2 µg DNase-treated RNA was reverse transcribed using a cDNA Synthesis Kit (Agilent Technologies, Santa Clara, CA, USA), following the respective manufacturer’s protocol. RT-PCR was performed in AriaMx (Agilent Technologies, software v1.0) using SYBR Green PCR Master Mix (iQ SYBR Green, BIO-RAD, CA, USA) (for primers, see **Table S7)**. The relative differences were determined by normalizing the Ct values of each gene to the mean expression of *β-Actin* and *Ubiquitin* genes and calculated using the equation 2^−ΔCt^ **(Livak and Schmittgen, 2001).**

### Extraction of volatile compounds and their identification

The volatile compounds from the fruits (5 biological replicates) were extracted as described by **Rambla et al. (2017).** The samples were processed as described in **Kilambi et al. (2021)**. The raw data were processed, the volatiles were identified and analysed as described in **Kilambi et al. (2021).**

### Statistical analysis

All results are expressed as mean±SE of three or more independent replicates. The StatisticalAnalysisOnMicrosoft-Excel (http://prime.psc.riken.jp-/MetabolomicsSoftware/StatisticalAnalysisOn-MicrosoftExcel/) was used to obtain significant differences between data points. Heat maps and 3D-PCA plots were generated using Morpheus (https://software.broadinstitute.org/morpheus/) and MetaboAnalyst-5.0 (https://www.metaboanalyst.ca/), respectively. The statistical significance was determined using Student’s t-test (for P≤0.05).

## Acknowledgments

This work was supported by the Department of Biotechnology (DBT), India grants, BT/PR11671/PBD/16/828/2008, BT/PR/7002/PBD/16/1009/2012, BT/COE/34/SP15209/2015 to RS and YS and BT/PR6983/PBD/16/1007/2012, BT/INF/22/SP44787/2021 to YS and RS. The Repository of Tomato Genomics Resources is a DBT-SAHAJ national facility.

## Author contributions

NRN, YS, and RS designed the research and wrote the manuscript; NRN designed genome editing, isolation, characterization, and RT-PCR of *rin^CR^* mutants; KS did biochemical characterization of mutants and crossed lines; PG helped in data analysis; IP and SS did RT-PCR of crossed lines. All authors read and approved the manuscript.

## Data Availability

The data consisting of genome editing, carotenoids, metabolites, and volatiles are included as supplemental material.

### Supplementary Data

**Table S1.** Identification of mutated alleles and zygosity in T_0_ transgenic plants.

**Table S2.** Mutation detection in potential off-target sites.

**Table S3.** The *Nps1* X *rin^CR^* (*rin^CR1^*and *rin^CR3^*) crossed F_2_ plants were examined for zygosity of *Nps1* and respective *rin^CR^* mutations at the seedling stage.

**Table S4.** On-vine monitoring of fruits of *Nps1* X *rin^CR1^* crossed F_2_ plants at different stages of fruit ripening.

**Table S5.** On-vine monitoring of fruits of *Nps1* X *rin^CR3^* crossed F_2_ plants at different stages of fruit ripening.

**Table S6.** List of primers used for RIN gRNA, plasmid construction, off-target monitoring, PCR-RE assay, and CAS9-free plant monitoring.

**Table S7.** List of primers used for quantitative real-time PCR of genes modulating ethylene biosynthesis, fruit ripening, carotenoids biosynthesis, and cell wall softening.

**Figure S1.** Schematic illustration of targeted *RIN* gene, plant transformation construct, and PCR-RE assay.

**Figure S2.** A representative image shows PCR-RE genotyping of T_2_ seedlings.

**Figure S3.** Comparison of on-vine-fruit ripening durations in *rin^CR1-4^* lines with AV fruits

**Figure S4.** Comparison of off-vine shelf-life in *rin^CR1-4^* lines with AV fruits.

**Figure S5.** The total amounts of carotenoids in *rin, rin^CR1-4^* mutants, and AV at different stages of ripening.

**Figure S6.** The distribution of carotenoids in *rin, rin^CR1-4^* mutants, and AV at RR stages of ripening.

**Figure S7.** Principle component analysis (PCA) of metabolites of *rin*, *rin^CR1-4,^* and AV fruits. **Figure S8.** The metabolic shifts in *rin* and *rin^CR1-4^* fruits during ripening compared to AV. **Figure S9.** The Brix and pH of AV, *rin,* and *rin^CR1-4^*mutants.

**Figure S10.** The Punnett square shows the phenotype of fruits of *Nps1* X *rin^CR1^* cross in F_1_ and F_2_ generations.

**Figure S11.** The Punnett square shows the phenotype of fruits of *Nps1* X *rin^CR3^* cross in F_1_ and F_2_ generations.

**Figure S12.** The total amounts of carotenoids in *rin^CR1^* and *rin^CR3^* mutants and the genetic segregation of carotenoids after crossing with the *Nps1* mutant.

**Figure S13.** The expression of ripening-related and cell-wall-related genes in *rin^CR1^* and *rin^CR3^* mutants and their genetic segregation after crossing with the *Nps1* mutant.

**Figure S14.** Comparison of on-vine fruit ripening in gene-edited *rin^CR1^* F_2_ lines generated by crossing with the *Nps1* mutant.

**Figure S15.** Comparison of on-vine fruit ripening in gene-edited *rin^CR3^* F_2_ lines generated by crossing with the *Nps1* mutant.

**Figure S16.** Comparison of post-harvest shelf-life in gene-edited *rin^CR1^* (and *rin^CR3^* lines crossed with *Nps1* mutant with AV.

**Figure S17.** Principle component analysis (PCA) of metabolites of *rin^CR1^* and *rin^CR3^* mutants (*n/n:r/r*) and their genetic segregation after crossing with the *Nps1* (mutant.

**Figure S18.** A comparison of genome-edited *RIN* alleles generated in different studies with the present study.

**Figure S19:** The expression of *RIN* and *phototropin1* genes in tomato fruits.

**Dataset S1**. The follow-up of edited RIN alleles from T_0_ till T_3_/T_4_ generation to obtain the CAS9-free homozygous edited plants.

**Dataset S2.** The levels of different carotenoids in *rin* mutant, *rin^CR^* lines, and AV at different stages of fruit ripening.

**Dataset S3.** The CO_2_ emission from *rin* mutant, *rin^CR^* lines, and AV at different stages of fruit ripening.

**Dataset S4.** The levels of different phytohormones in *rin* mutant, *rin^CR^* lines, and AV at different stages of fruit ripening.

**Dataset S5.** The levels of different metabolites in *rin* mutant, *rin^CR^* lines, and AV at different stages of fruit ripening.

**Dataset S6.** The expression of ripening-related and cell-wall-related genes in AV, *rin,* and

*rin^CR1-4^* fruits at different stages of ripening.

**Dataset S7.** The levels of different carotenoids in RR fruits *rin^CR1^* and ri*n^CR3^* lines crossed with *Nps1* in the F_2_ generation.

**Dataset S8**. The expression of ripening-related and cell-wall-related genes in RR fruits of *rin^CR1^* and ri*n^CR3^* lines crossed with *Nps1* in the F_2_ generation.

**Dataset S9.** The levels of different phytohormones in RR fruits of *rin^CR1^* and ri*n^CR3^* lines crossed with *Nps1* in the F_2_ generation.

**Dataset S10**. The levels of different metabolites in RR fruits of *rin^CR1^* and ri*n^CR3^* lines crossed with *Nps1* in F_2_ generation.

**Dataset S11.** The levels of different volatiles in RR fruits of *rin^CR1^* and ri*n^CR3^* lines crossed with *Nps1* in F_2_ generation.

## Abbreviations

AV: - Arka Vikas,
*rin*: - ripening inhibitor.

## Legends to Supplementary Figures

**Figure S1:**
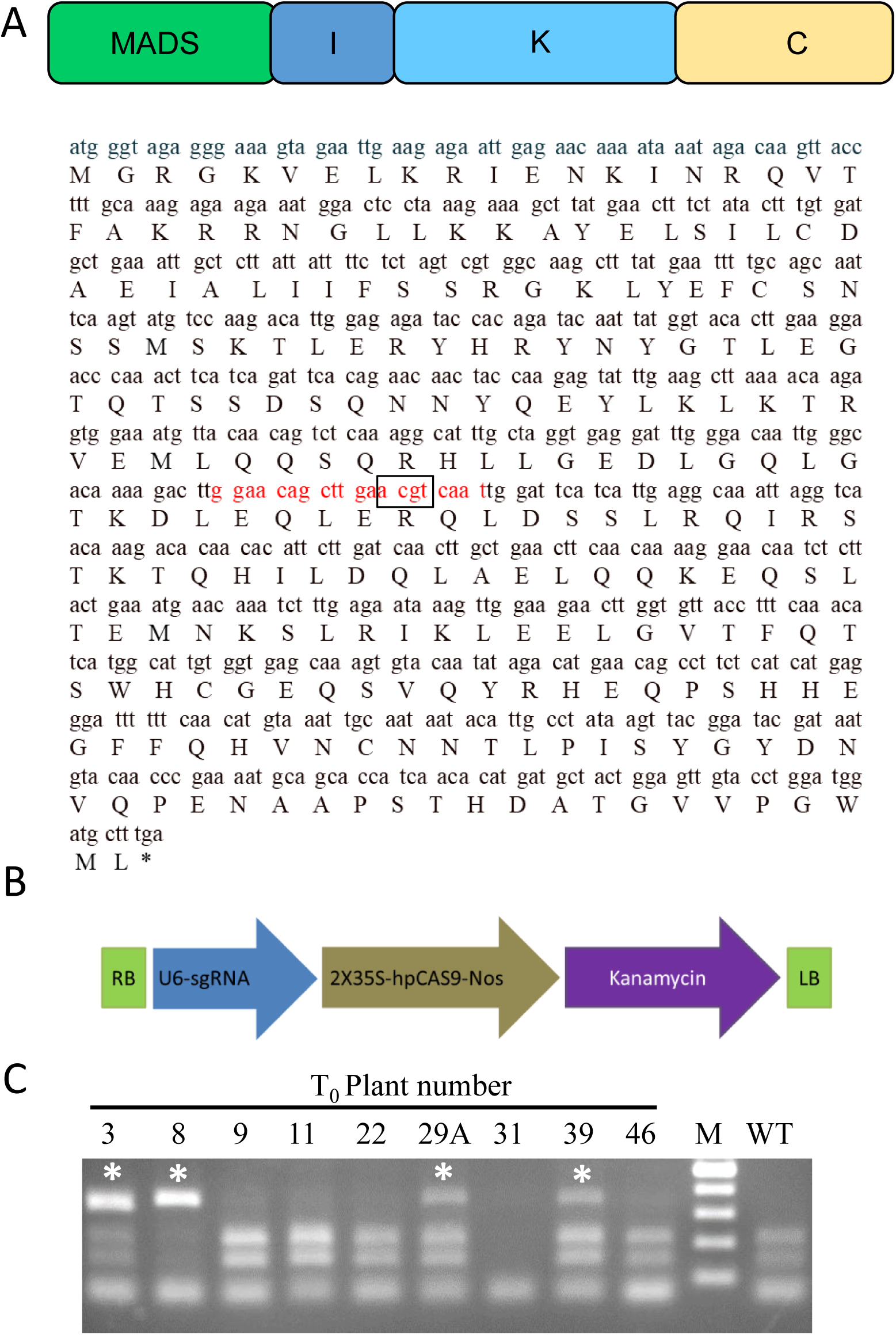
Schematic illustration of targeted RIN gene, plant transformation construct, and PCR-RE assay. **A.** The domain structure, cDNA, and protein sequence of RIN MADS-box transcription factor (NCBI Reference Sequence: NM_001247741.2; Wang et al., 2020). Red-colored letters indicate the 20 bp sequence targeted by gRNA. The TaiI restriction site ACGT^ is enclosed in a rectangle. **B**. Schematic representation of binary vector used in this study. U6-sgRNA: Arabidopsis *U6* promoter and the gRNA sequence, 2X35S-hpCAS9-Nos: 2X*CaMV35S* promoter sequence, hpCas9: human-codon optimized SpCas9, Nos: Nos terminator, Kanamycin: the kanamycin-resistant marker expression cassette, RB: right border of T-DNA, LB: left border of T-DNA. **C**. A representative image shows the screening of *RIN* gene knockout induced by the CRISPR/Cas9 system using PCR-RE analysis of the target site. The target region was PCR amplified from genomic DNA extracted from transgenic plants and then digested with TaiI restriction enzyme. A white asterisk in the lane indicates the loss of the restriction site targeted by the TaiI enzyme. Plants 3, 29A, and 39 are heterozygous for editing, while plant 8 is homozygous. **M:** marker; **WT:** wild-type control

**Figure S2:**
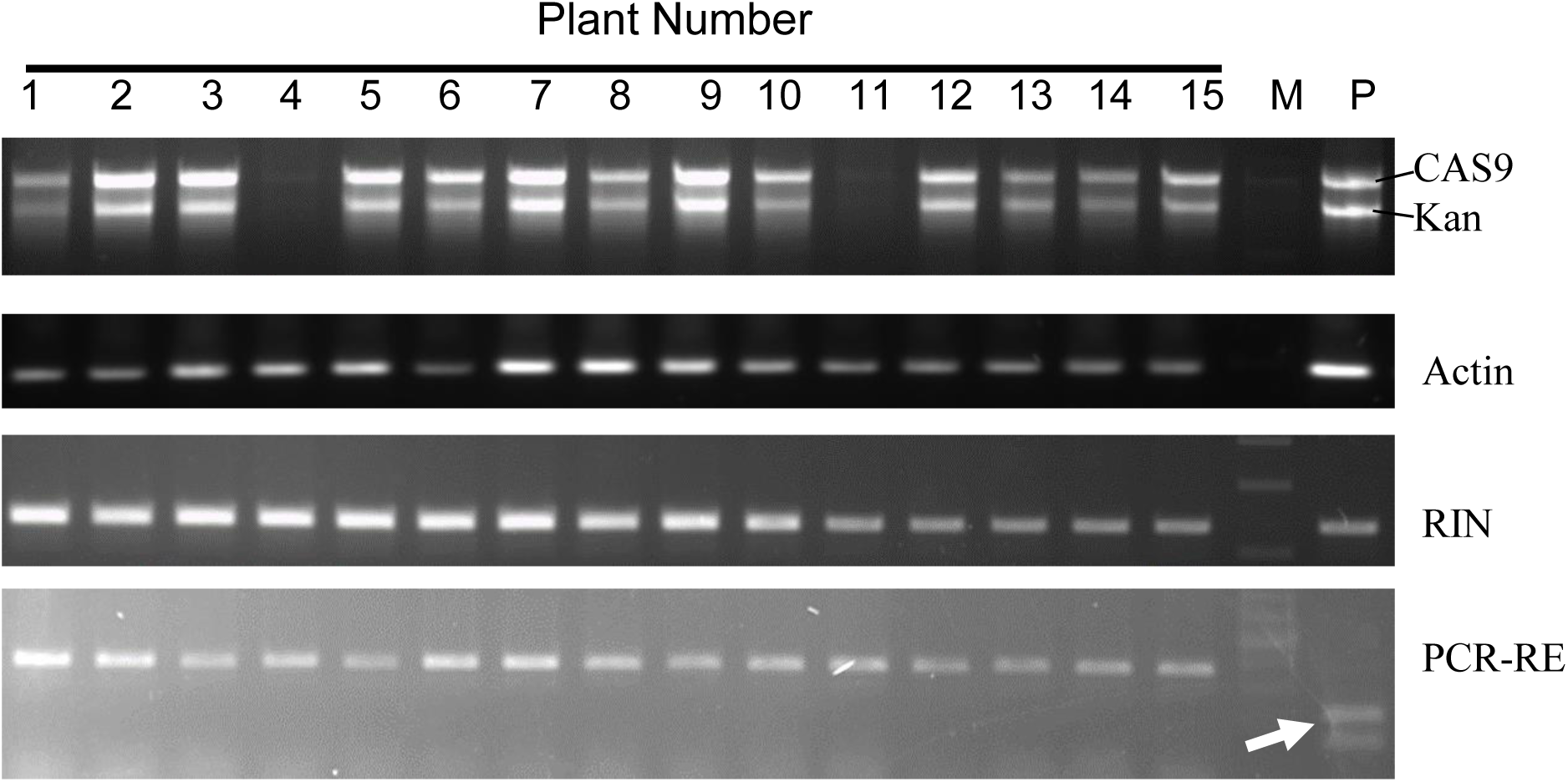
A representative image shows PCR-RE genotyping of T_2_ seedlings. The gRNA target region of *RIN* was *PCR* amplified from genomic DNAs extracted from T_2_ seedlings. Housekeeping gene actin was used as an internal control for the PCR genotyping. Post-PCR amplification, the product was subjected to PCR-RE assay with TaiI enzyme. The absence of the TaiI restriction site in the PCR product indicates that the plant was homozygous for *RIN* editing in the T_1_ generation. Plants 4 and 11 are CAS9-free and Kanamycin (Kan) free. Lanes 1-15: T_2_ progeny plants of *rin^CR1^*. **M:** marker; **P:** respective positive controls. The white arrow indicates wild-type cleavage of PCR product on TaiI digestion.

**Figure S3.**
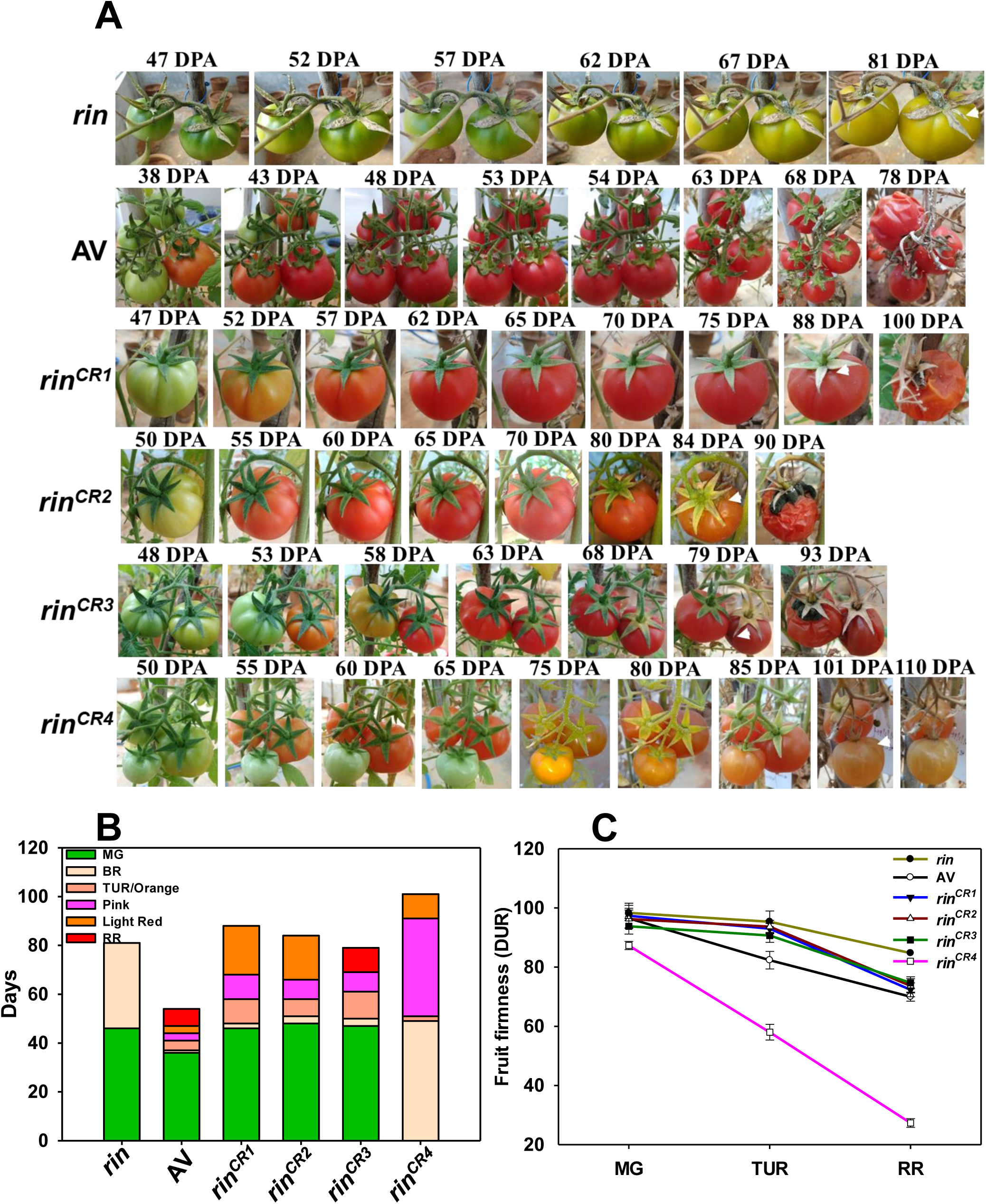
Comparison of on vine-fruit ripening durations in gene-edited *rin^CR1-4^* lines with Arka Vikas (AV) fruits. A. On-vine ripening of AV, *rin*, and *rin^CR1-4^* mutants in T_3_ generation. The photographs show the ripening progression until the onset of the senescence (White arrowheads). The appearance of wrinkles on fruit skin was considered as the time point of senescence onset [AV (54 DPA), *rin* (81 DPA), *rin^CR1^* (88 DPA), *rin^CR2^* (84 DPA), *rin^CR3^* (79 DPA), and *rin^CR4^* (101 DPA)] Note that unlike AV (RR at 43 DPA), the *rin^CR1-4^* mutants do no attain RR stage, but reach light red (*rin^CR1-3^* or pink stage (*rin^CR4^*). The *rin* mutant, though, shows the loss of green color but does not show coloration like AV or *rin^CR1-4^* mutants. B. Chronological development of tomato fruits post-anthesis until the onset of senescence (n = 3). The stacked bar graph shows the attainment of different stages of ripening: mature green (MG), breaker (BR), turning (TUR), pink (P), and red ripe (RR). C. Fruit firmness of AV, *rin*, and *rin^CR1-4^* mutants. The fruit firmness was measured after harvesting the fruits at MG, TUR, and RR stages of ripening. The firmness value was recorded by measuring each fruit two-three times at the equatorial plane. The *rin^CR4^* fruits were very soft compared to AV and other *rin* alleles. Even being soft, they developed wrinkling of skin similar to other *rin^CR1-3^* fruits > 90 dpa. (mean ± SE, n = 5).

**Figure S4.**
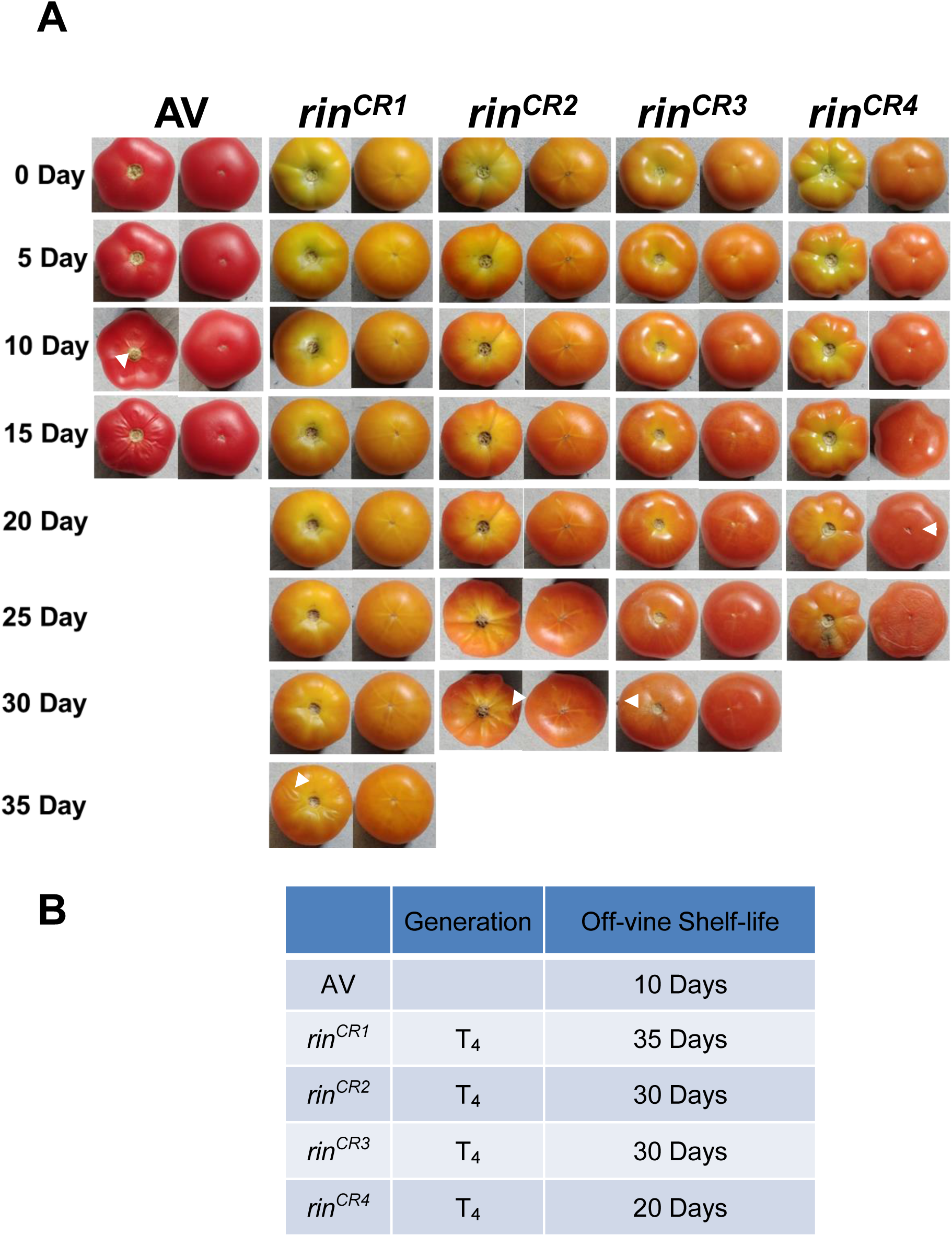
Comparison of off-vine shelf-life in gene-edited *rin^CR1-4^* lines with Arka Vikas (AV) fruits. A. The fruits were harvested at the RR stage for AV and the pink stage for *rin^CR1-4^* mutants. The photographs were taken at 5-day intervals until the appearance of wrinkles on the fruit surface (White arrowheads). The appearance of wrinkles was taken as the time point of senescence onset. Like on-vine ripening, the *rin^CR4^* fruits were softer and developed wrinkles much earlier than *rin^CR1-3^.* B. The table shows the time points of wrinkle appearance in *rin^CR1-4^* lines.

**Figure S5:**
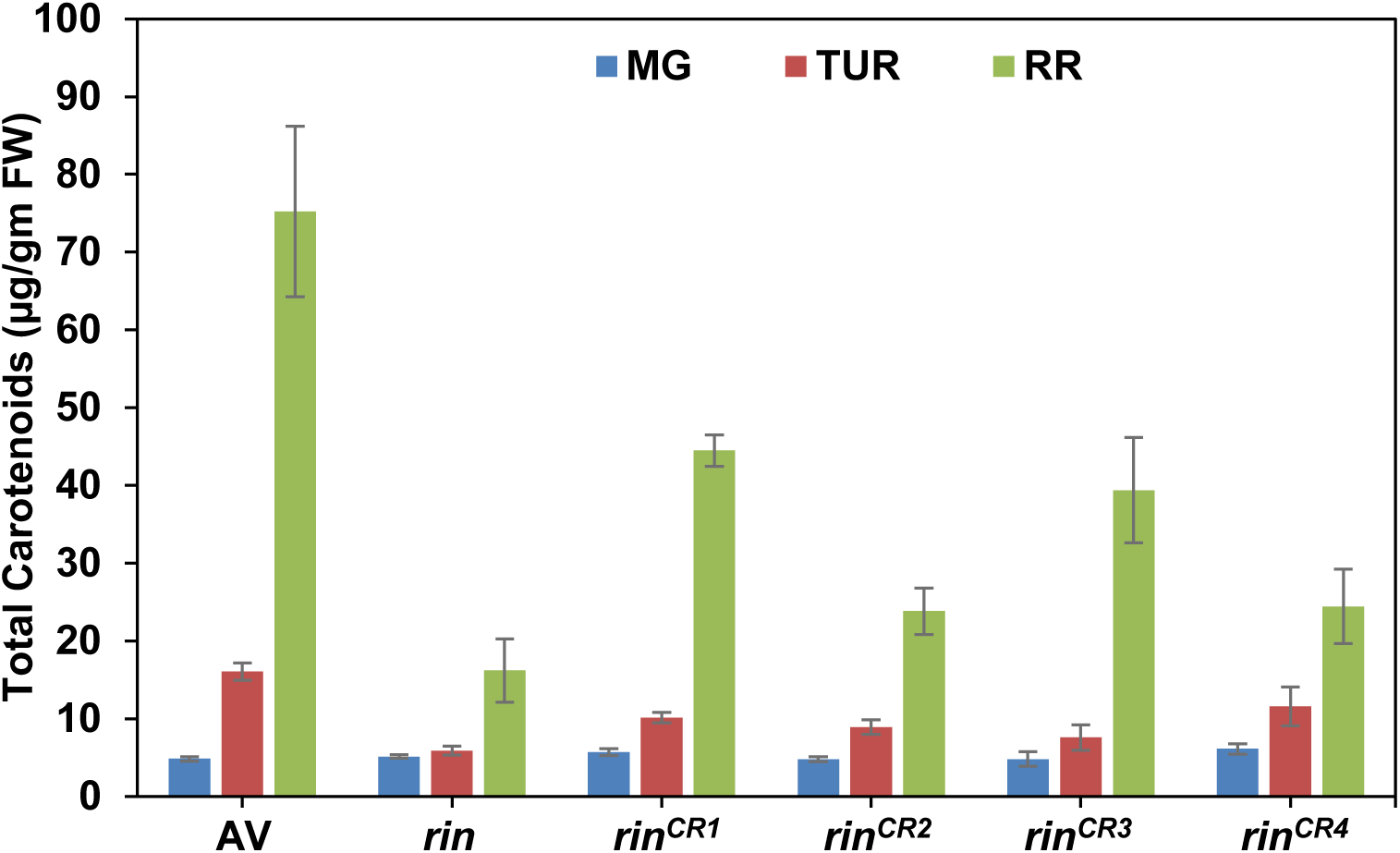
The total amounts of carotenoids in *rin, rin^CR1-4^* mutants, and Arka Vikas (AV) at different stages of ripening. Unlike *rin^CR1-4^* alleles, the *rin* mutant does not accumulate phytoene, phytofluene, and lycopene during ripening. At the RR stage, the *rin^CR1-4^* mutants have 28-54% carotenoids compared to AV. The carotenoid data are means ± SE (n ≥ 3) *P ≤ 0.05. See **Dataset S2** for individual carotenoid levels and significance.

**Figure S6:**
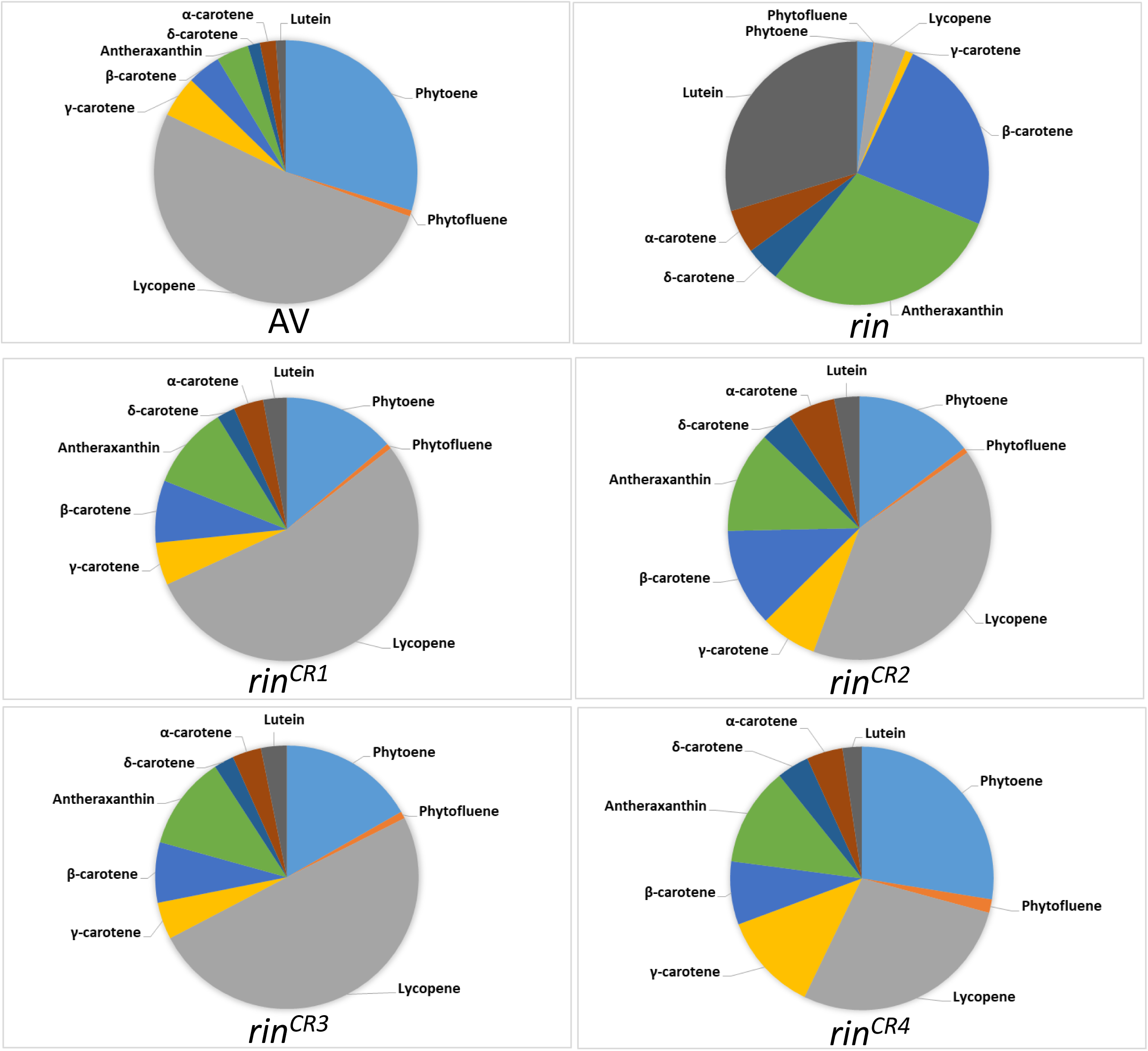
The distribution of carotenoids in *rin, rin^CR1-4^* mutants, and Arka Vikas (AV) at RR stages of ripening. The pie diagram depicts individual carotenoids, as a percent of the total carotenoids (100%) present, in wild type, in *rin*, and in each *rin^CR^* allele. Note the carotenoid distribution in the *rin^CR^* alleles is somewhat intermediate between *rin* and AV. See **Dataset S2** for individual carotenoid percentages.

**Figure S7.**
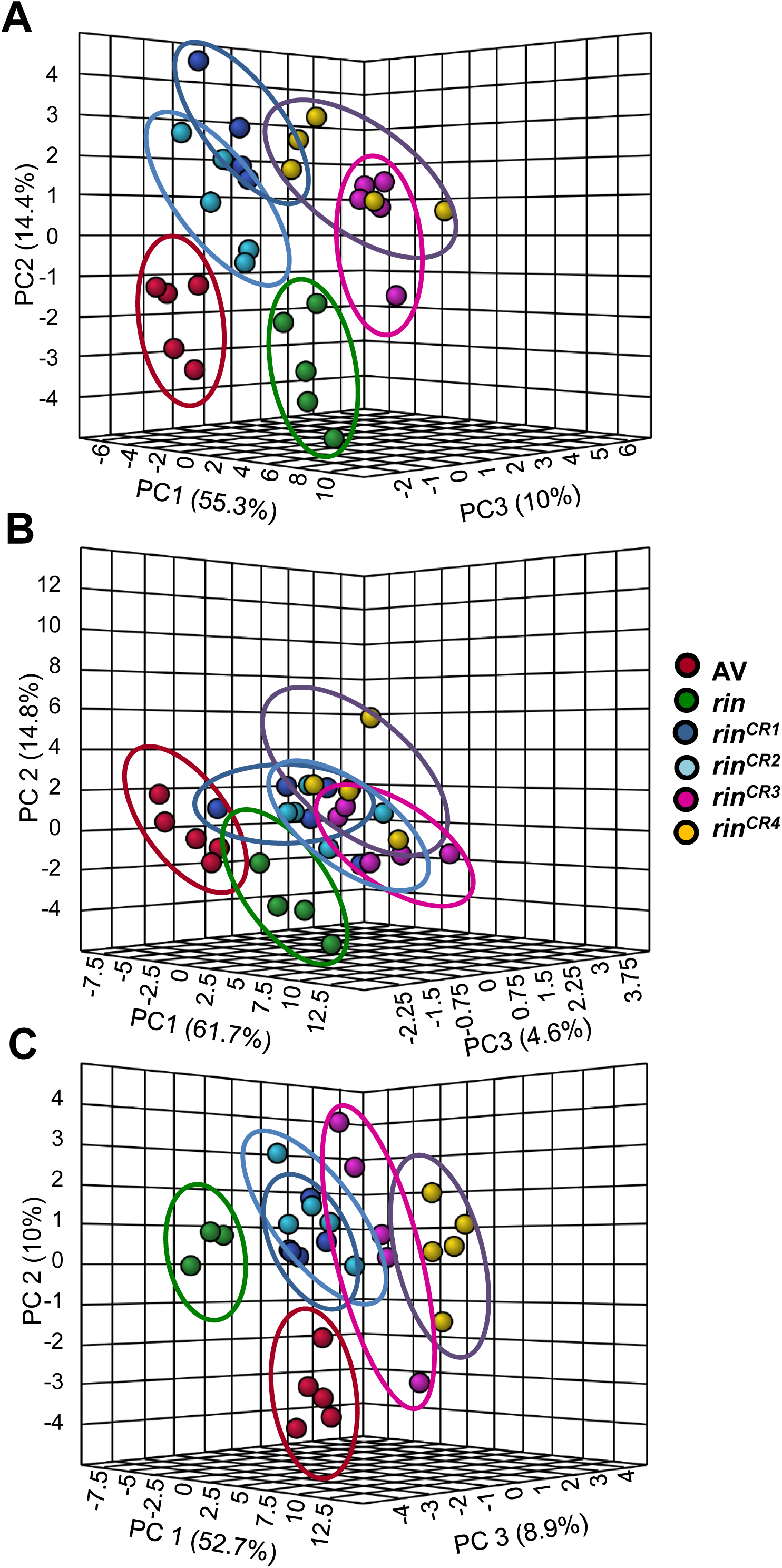
Principle component analysis (PCA) of metabolites of *rin*, *rin^CR1-4^*, and Arka Vikas (AV) fruits. (**A**) MG, **(B**) TUR, (**C**) RR. The PCA was constructed using MetaboAnalyst 5.0. The variance of the PC1, PC2, and PC3 is given in parentheses.

**Figure S8.**
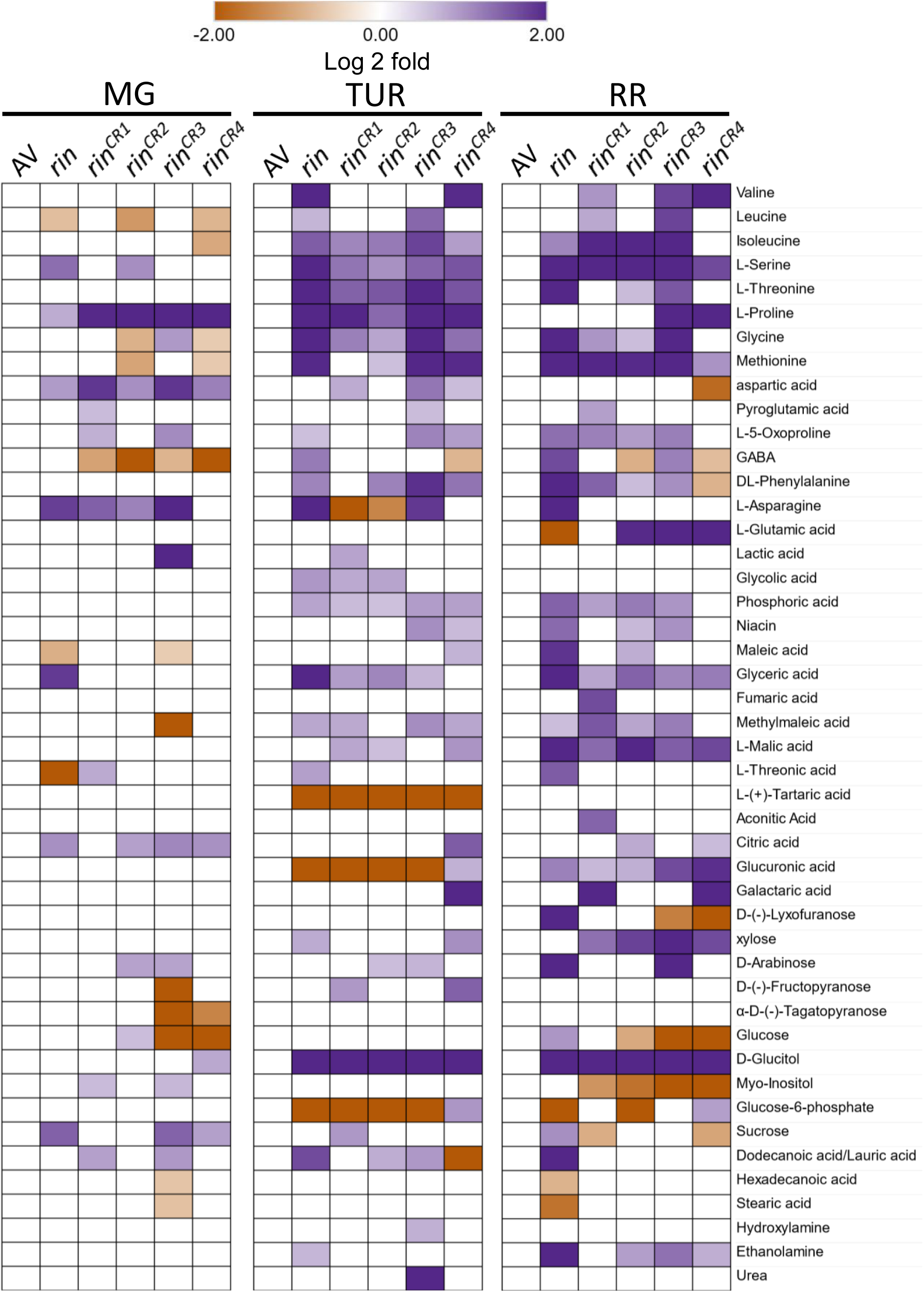
The metabolic shifts in *rin* and *rin^CR1-4^* fruits during ripening compared to Arka Vikas (AV). The relative changes in the metabolite levels at MG, TUR, and RR stages were determined by calculating the log2 fold change at respective ripening phases with reference to AV. Only significantly changed metabolites are depicted (Log2 fold ≥ ± 0.584, p-value ≤ 0.05, n ≥ 5)). See **Dataset S5** for detailed metabolite data.

**Figure S9.**
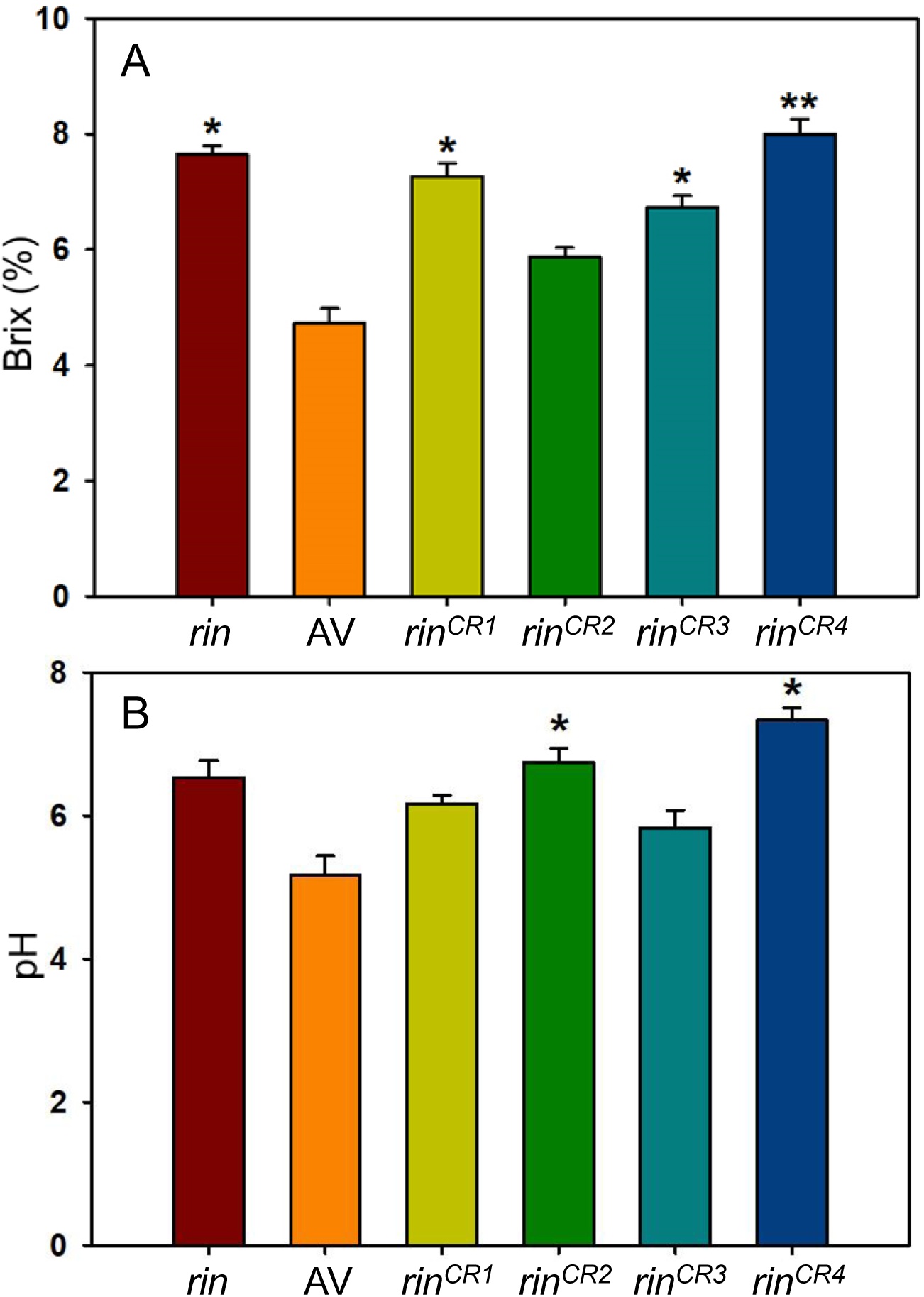
Fruit total soluble solids (°Brix) (A) and pH (B) of AV, *rin*, and *rin^CR1-4^* mutants. (Student’s t-test; * for P ≤ 0.05, ** for P ≤ 0.01 and *** for P ≤ 0.001, n=3 ± SE).

**Figure S10.**
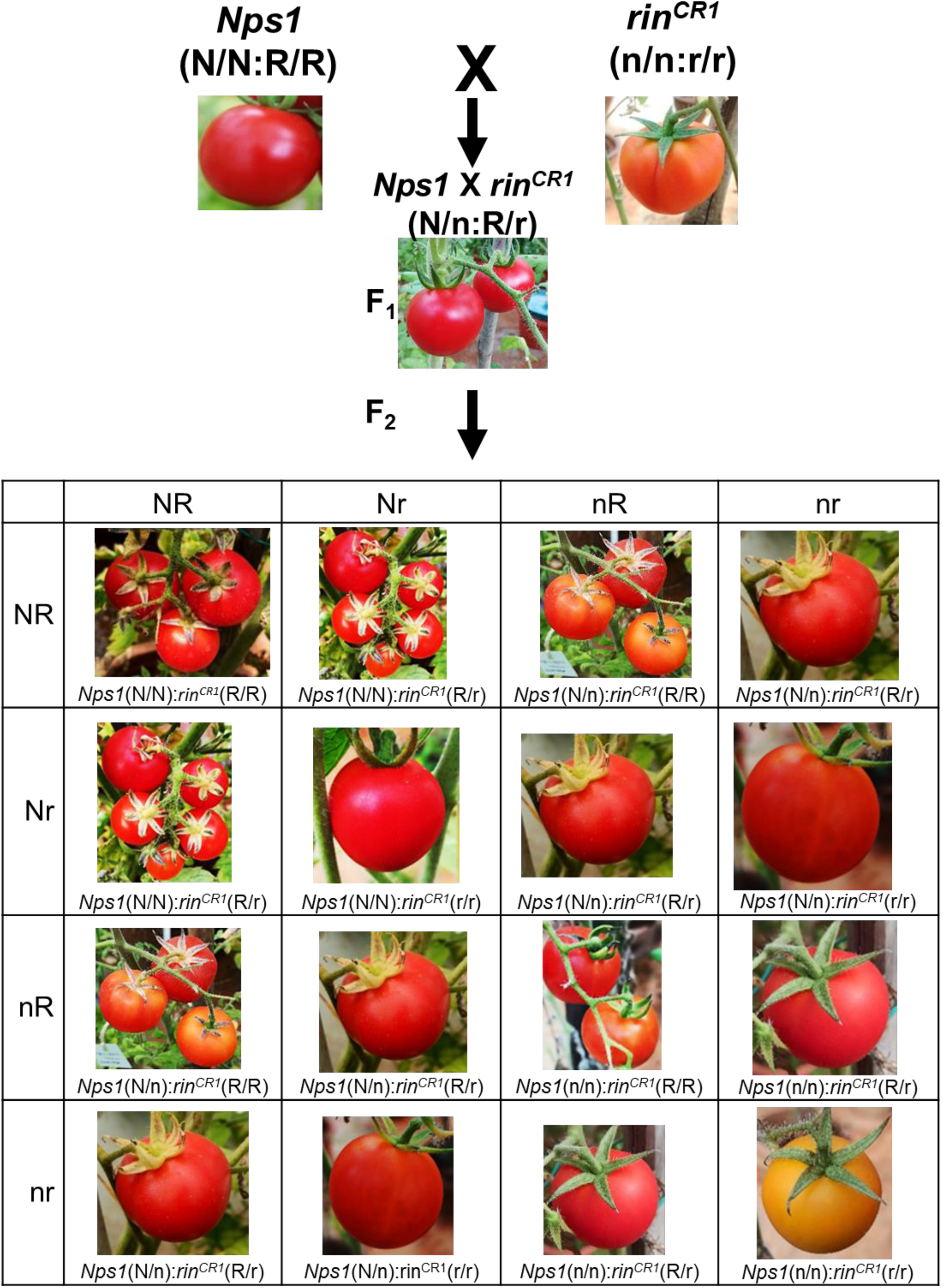
The Punnett square shows the phenotype of fruits of *Nps1* X *rin^CR1^* cross in F_1_ and F_2_ generations. *Nps1* being the dominant gene, the homozygous (*N/N*) or heterozygous (*N/n*) fruits accumulate a higher amount of carotenoids, suppressing *rin^CR1^* influence on the reduction of carotenoids. Note light-red color fruit phenotype of *rin^CR1^* reappears in the *n/n:r/r* combination. For the segregation ratio, see Table S3.

**Figure S11.**
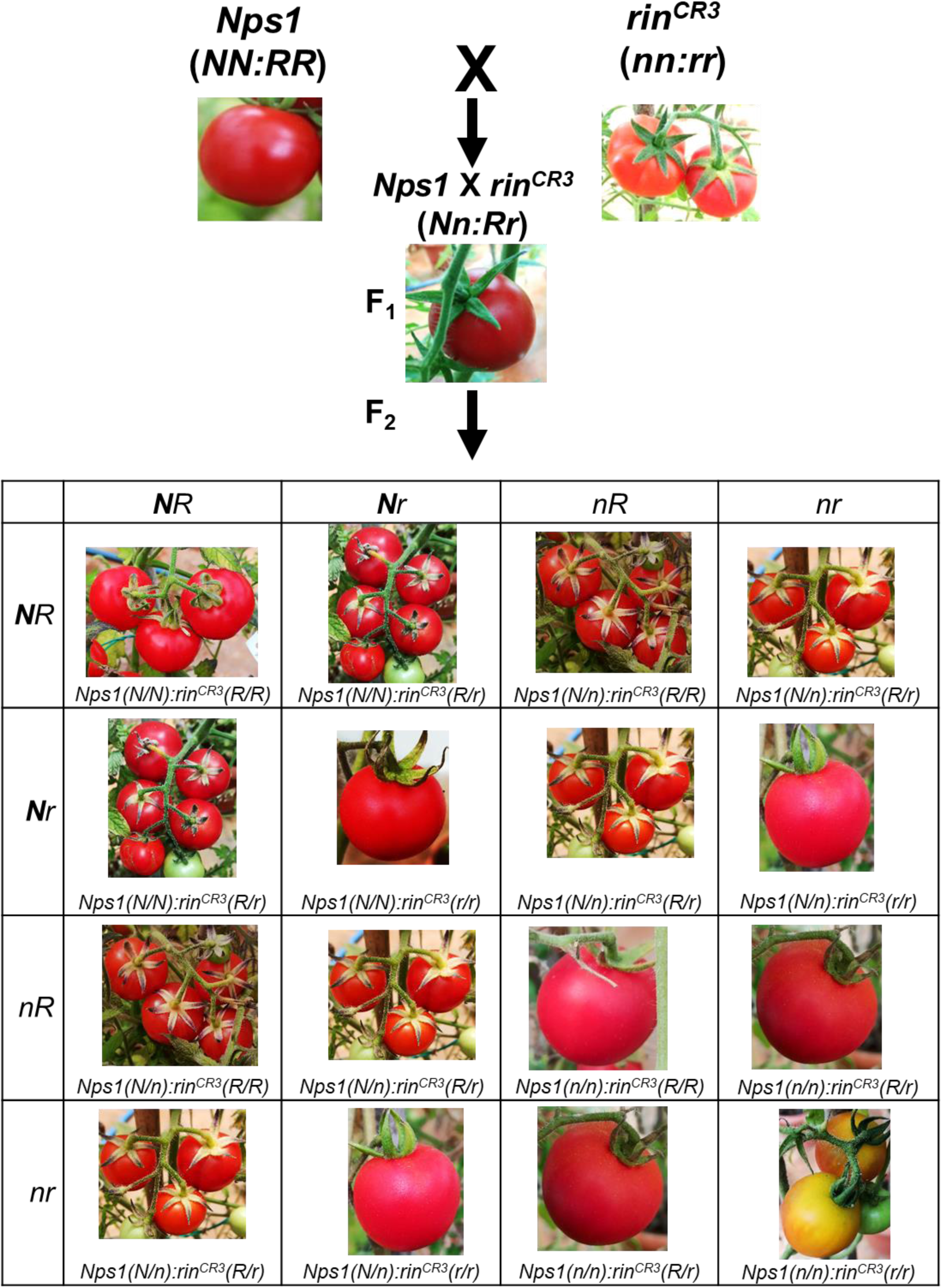
The Punnett square shows the phenotype of fruits of *Nps1* X *rin^CR3^* cross in F_1_ and F_2_ generations. *Nps1* being the dominant gene, the homozygous (*N/N*) or heterozygous (*N/n*) fruits accumulate a higher amount of carotenoids, suppressing *rin^CR3^* influence on the reduction of carotenoids. Note light-red color fruit phenotype of *rin^CR3^* reappears in the *n/n:r/r* combination. For the segregation ratio, see Table S3.

**Figure S12:**
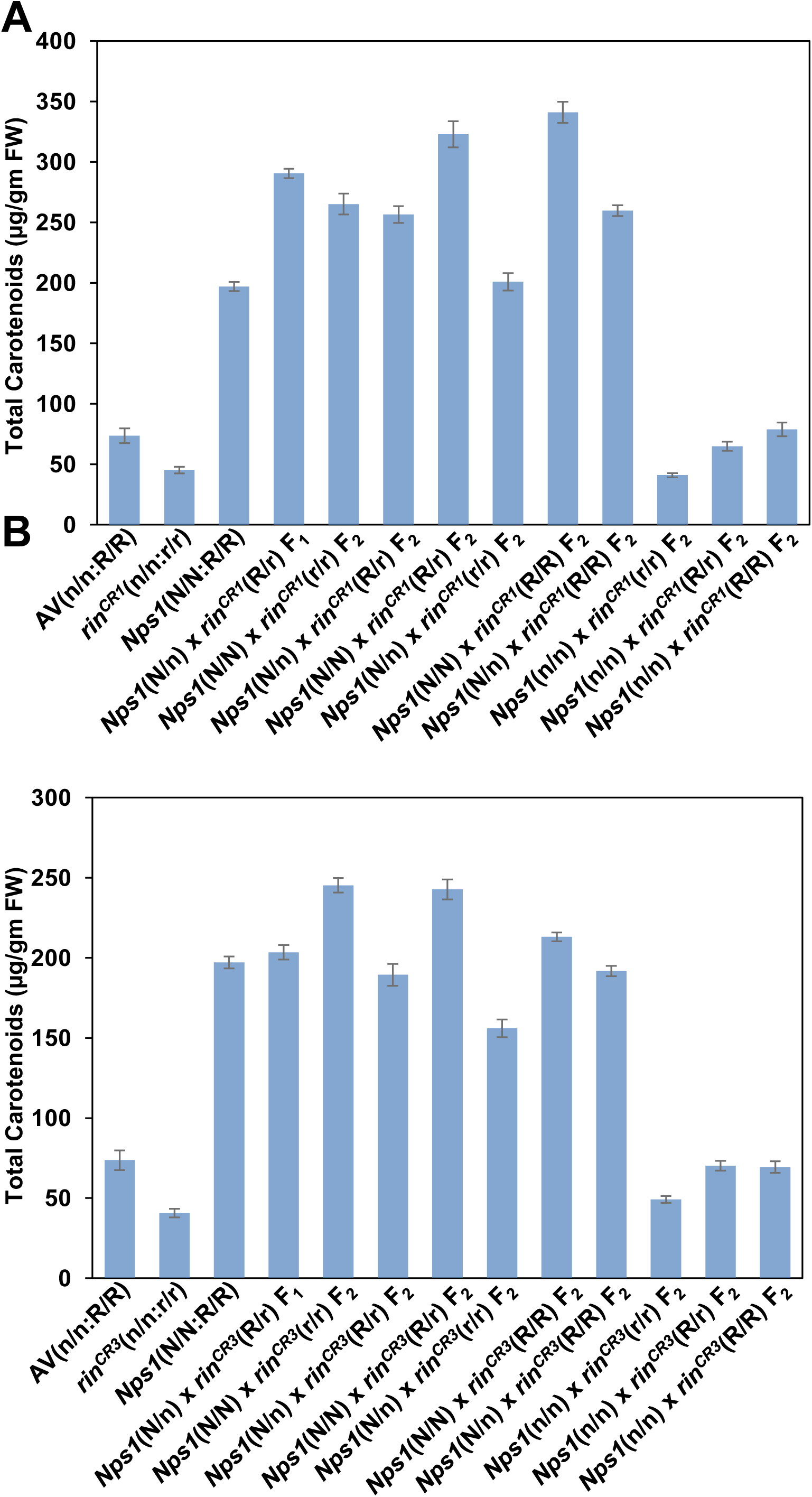
The total amounts of carotenoids in *rin^CR1^* (**A**) and *rin^CR3^* (**B**) mutants (*n/n:r/r*) and the genetic segregation of carotenoids after crossing with the *Nps1* (*N/N:R/R*) mutant. The carotenoids were analysed at the RR stage. Note that *Nps1* being a dominant mutation, enhances carotenoids in homozygous (*N/N*) and also heterozygous (*N/n*) states in *rin^CR1^* and *rin^CR3^* alleles and in wild-type (*R/R*). The homozygous F_2_ lines bearing the wild-type allele (*n/n:R/R*) have carotenoids like Arka Vikas (AV) (*n/n:R/R*), and the one bearing the *rin^CR1^* or *rin^CR3^* (*n/n:r/r*) is like their respective allele. The carotenoid data are means ± SE (n ≥ 3) *P ≤ 0.05. See **Dataset S7** for individual carotenoid levels and significance.

**Figure S13:**
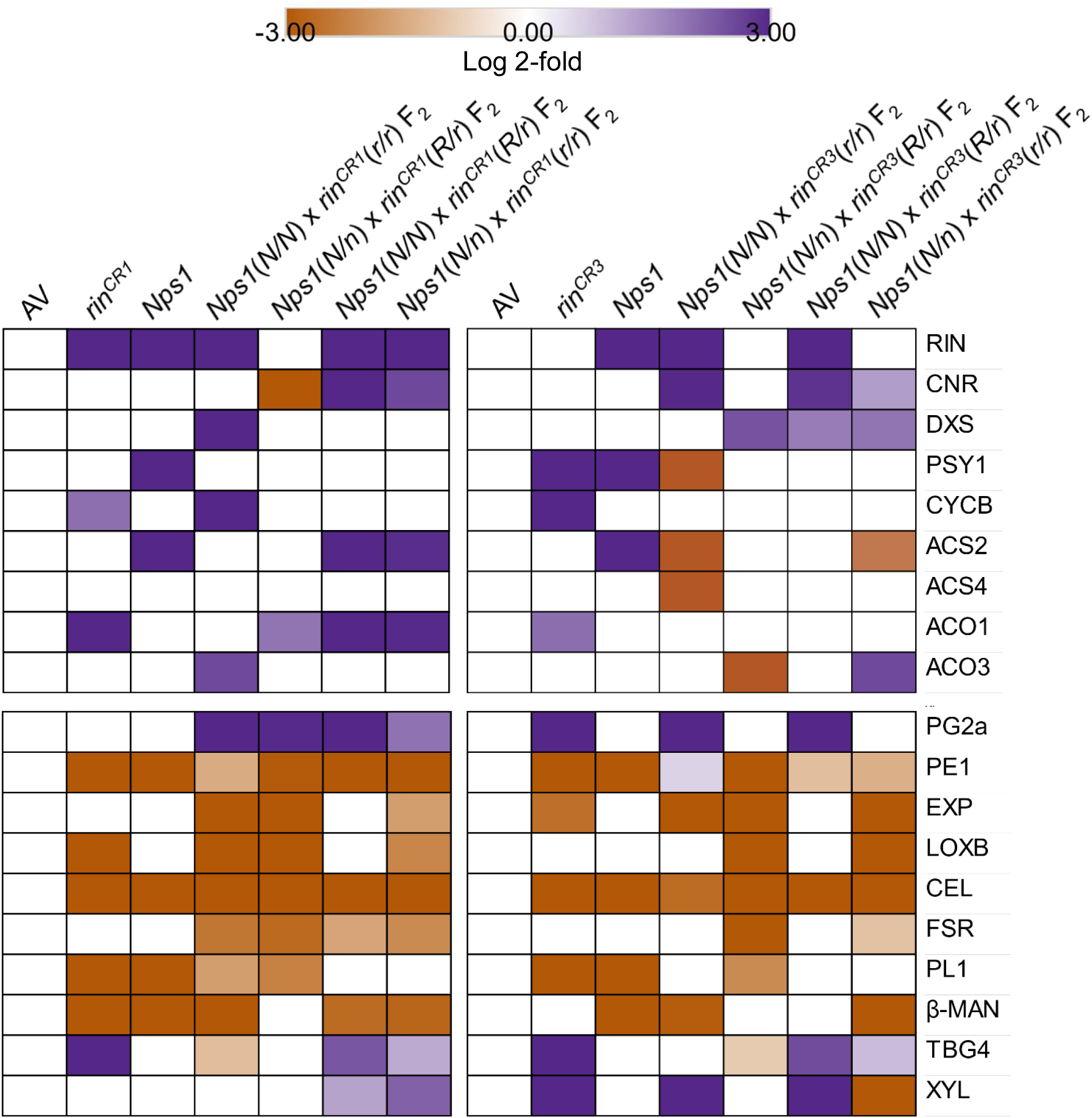
The expression of ripening-related (**Upper panel**) and cell-wall-related (**Lower panel**) genes *rin^CR1^* and *rin^CR3^* mutants (*n/n:r/r*) and their genetic segregation after crossing with the *Nps1* (*N/N:R/R*) mutant. The gene expressions were analysed at the RR stage. The relative changes in gene expression were determined by calculating the log2 fold change with reference to Arka Vikas (AV). Only significantly changed transcripts are depicted (Log2 fold ≥ ± 0.584, p-value ≤ 0.05, n ≥ 3)). See **Dataset S8** for detailed gene expression data. *Gene abbreviation*: *ACS2* -1-amino-cyclopropane-1-carboxylate synthase 2, *ACS4* -1-amino-cyclopropane-1-carboxylate synthase4, *ACO1* -1-amino-cyclopropane-1-carboxylate oxidase1, *ACO3* -1-amino-cyclopropane-1-carboxylate oxidase3, *RIN* - Ripening inhibitor, *CNR* -colorless non-ripening, *DXS* -1-deoxy-D-xylulose 5-phosphate synthase, *PSY1* -Phytoene synthase1, *CYCB* -Lycopene β-cyclase, *CEL* -Cellulase, *EXP1* -Expansin 1, *PG2a* - Polygalacturonase 2A, *PE1* -Pectin esterase 1, *FSR* -Fruit shelf-life regulator, *PL* -pectate lyase, *β-MAN* -β-mannosidase, *TBG4* -tomato β-galactosidase 4, *XYL* -β-D-xylosidase, *LOXB* -Lipoxygenase B.

**Figure S14.**
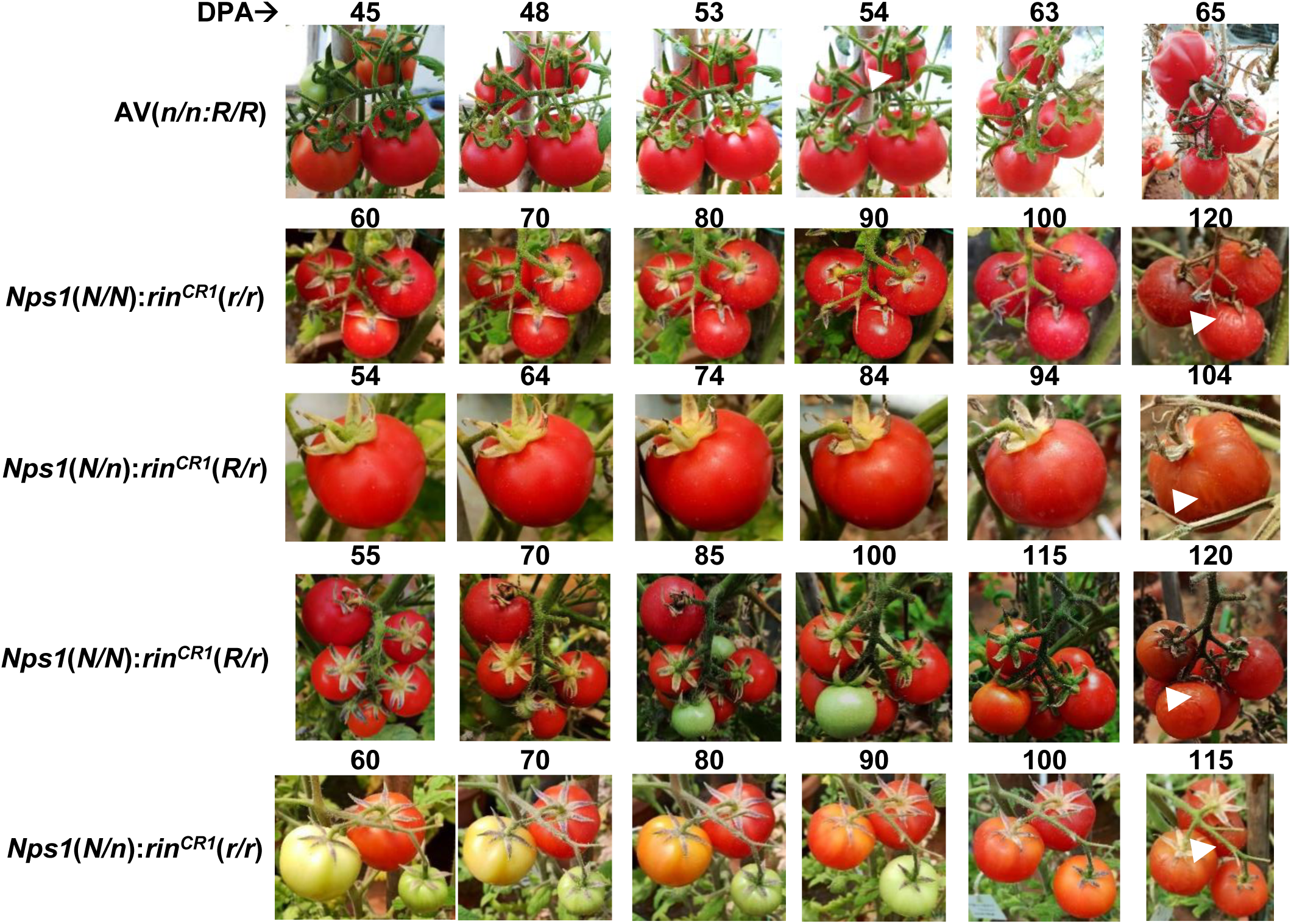
Comparison of on-vine fruit ripening in gene-edited *rin^CR1^* (*n/n:r/r*) F_2_ lines generated by crossing with the *Nps1* (*N/N:R/R*) mutant. Photographs show the on-vine ripening of four *Nps1/rin^CR1^* crosses, where both alleles are present in either homozygous or heterozygous conditions. The ripening progression was monitored from the red-ripe stage till the onset of the senescence (White arrowheads). The appearance of wrinkles on fruit skin was considered as the time point for senescence onset. Note that compared to Arka Vikas (AV), the dihybrid fruits show an extended on-vine red-ripe stage. For details of the ripening stages, see Table S4.

**Figure S15.**
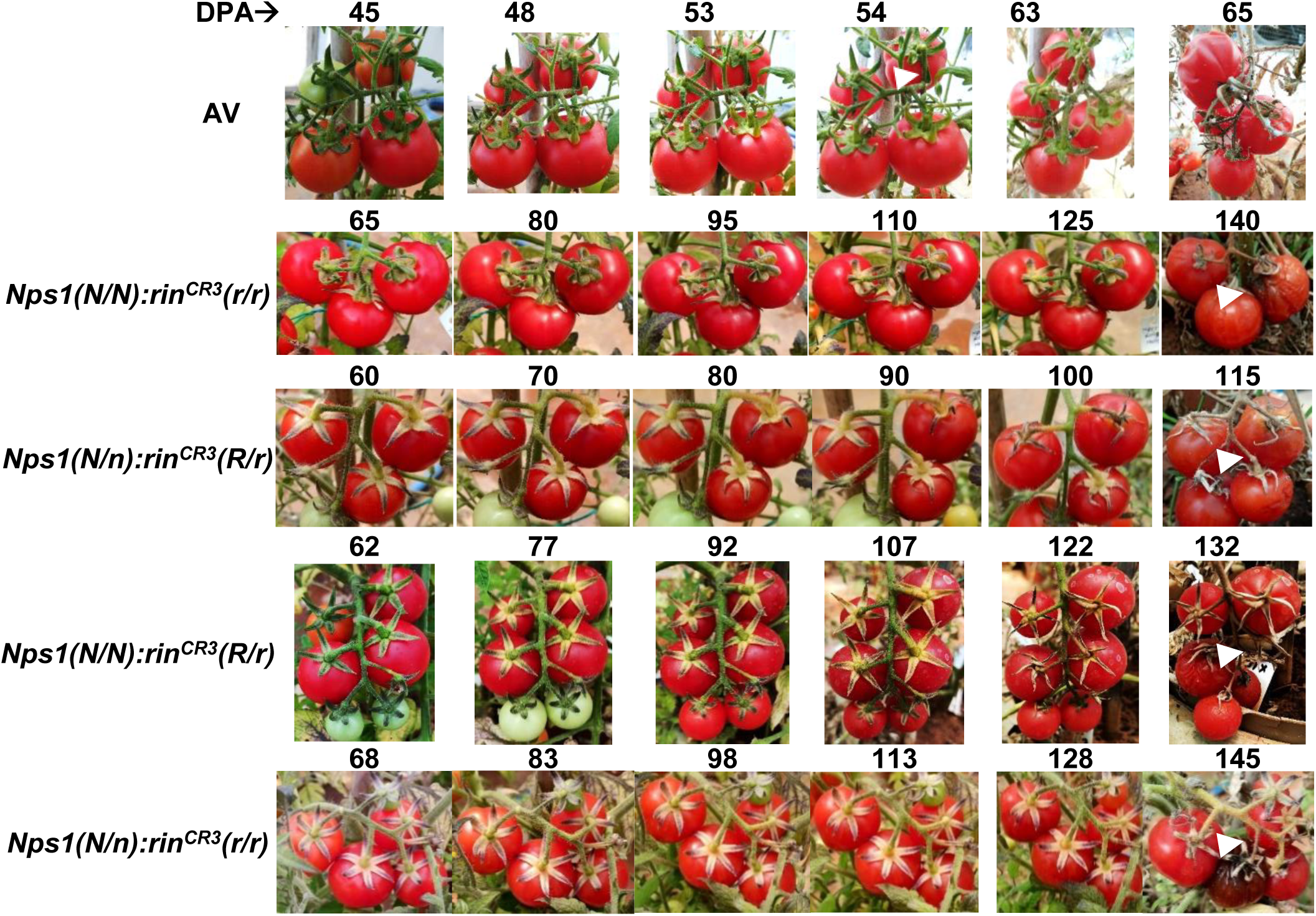
Comparison of on-vine fruit ripening in gene-edited *rin^CR3^* (*n/n:r/r*) F_2_ lines generated by crossing with the *Nps1* (*N/N:R/R*) mutant. Photographs show the on-vine ripening of four *Nps1/rin^CR3^* crosses, where both alleles are present in either homozygous or heterozygous conditions. The ripening progression was monitored from the red-ripe stage till the onset of the senescence (White arrowheads). The appearance of wrinkles on fruit skin was taken as the time point for senescence onset. Note that compared to Arka Vikas (AV), the dihybrid fruits show an extended on-vine red-ripe stage. For details of the ripening stages, see Table S5.

**Figure S16.**
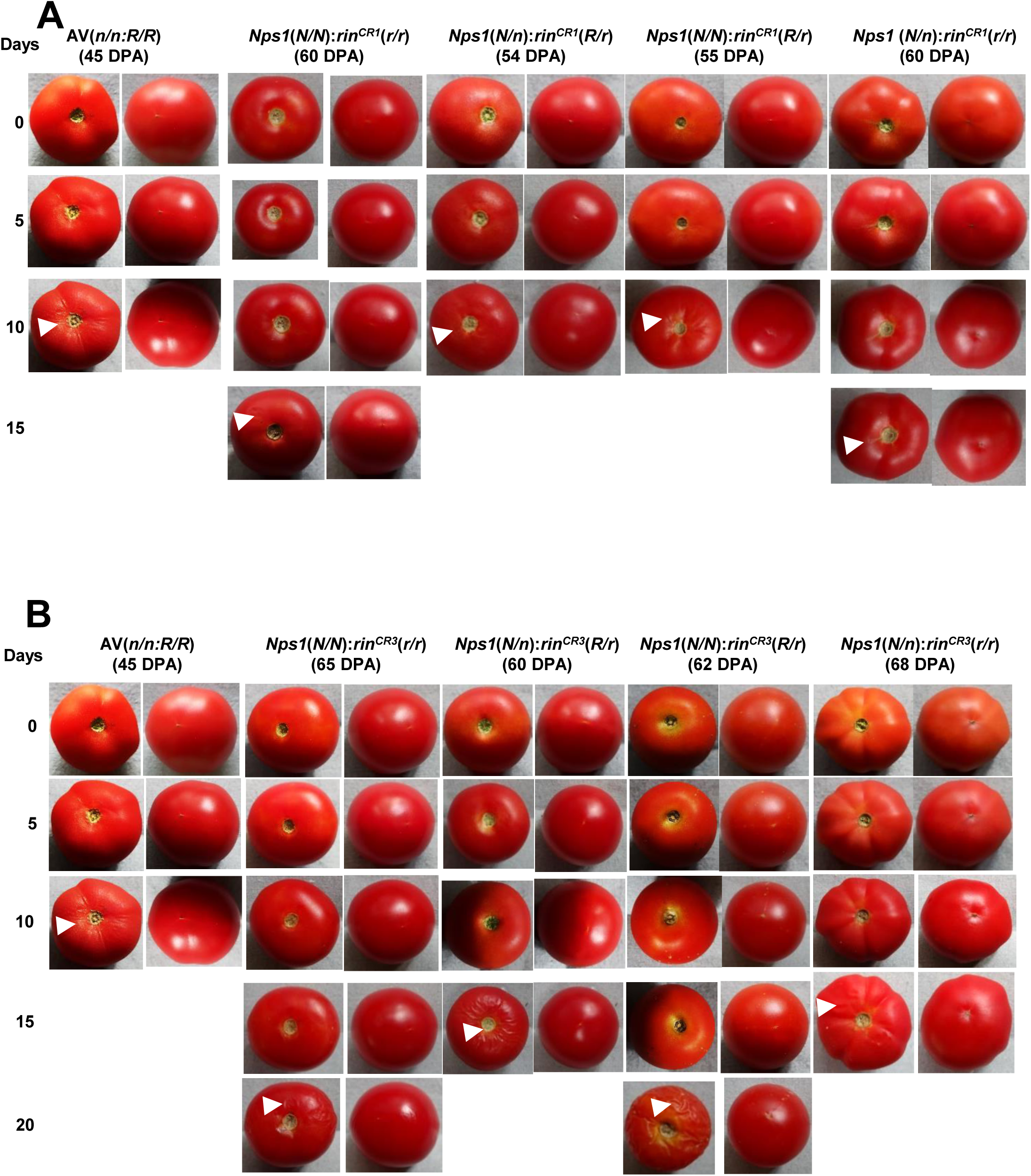
Comparison of post-harvest shelf-life in gene-edited *rin^CR1^* (A) and *rin^CR3^* (B) lines (*n/n:r/r*) crossed with *Nps1* mutant (*N/N:R/R*) with AV (*n/n:R/R*). The fruits were harvested at the RR stage for AV and *rin^CR^* mutants. The photographs were taken at 5-day intervals until the appearance of wrinkles on the fruit surface (White arrowheads). The appearance of wrinkles was taken as the time point of senescence onset. Among the two *rin^CR^* alleles, the fruits of the *rin^CR3^* allele, having RIN protein truncation, display longer off-vine shelf life. The representative data from three different biological replicates are presented.

**Figure S17.**
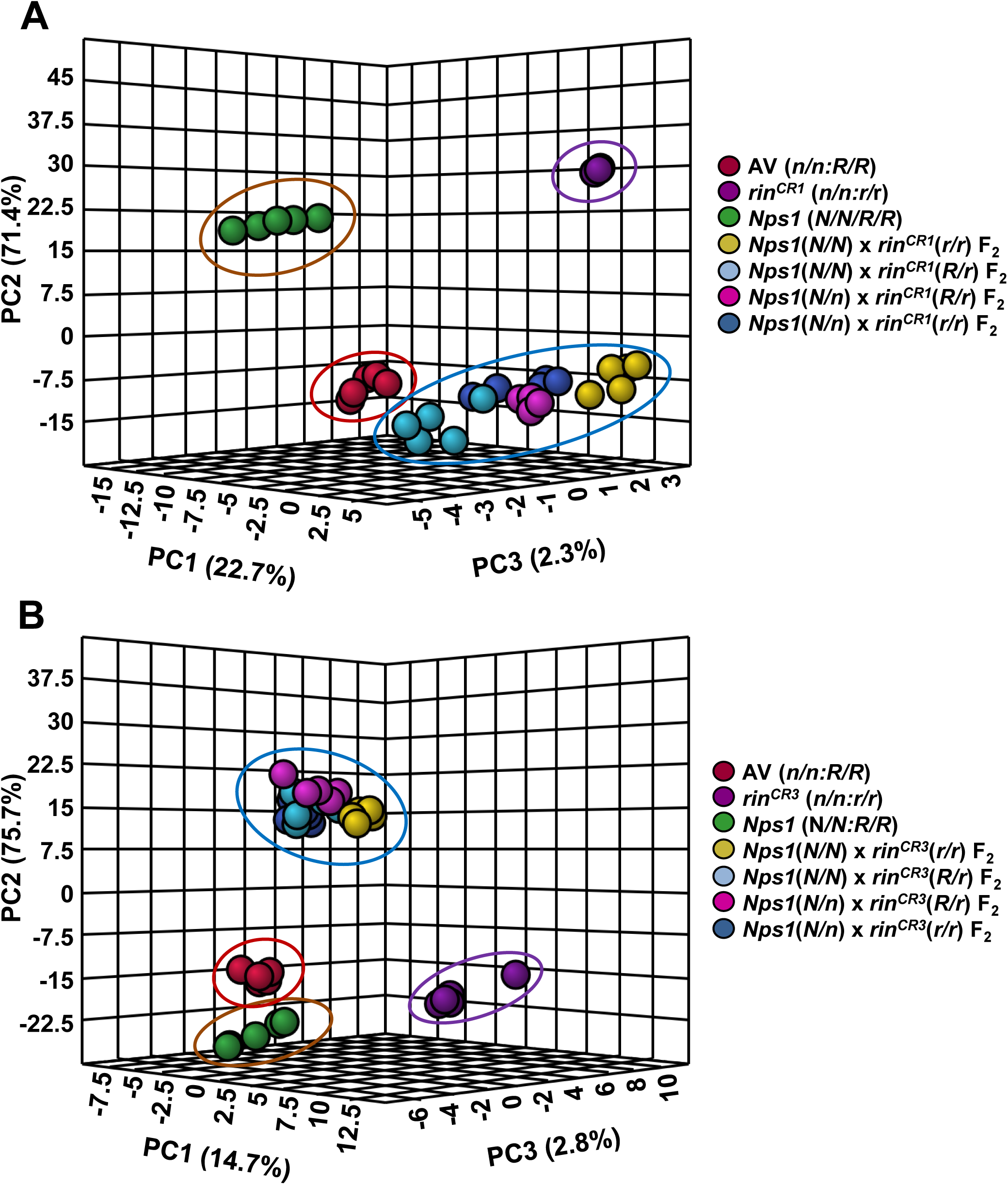
Principle component analysis (PCA) of metabolites of *rin^CR1^* (**A**) and *rin^CR3^* (**B**) mutants (*n/n:r/r*) and their genetic segregation after crossing with the *Nps1* (*N/N:R/R*) mutant. The metabolites were analysed at the RR stage. Panel **A** and **B** show PCA of four *Nps1/rin^CR1^* and four *Nps1/rin^CR3^* F_2_ crosses, where both the alleles are present in either homozygous or heterozygous conditions. Note that PCAs of all four F_2_ combinations of *Nps1/rin^CR1^* and *Nps1/rin^CR3^* are in close proximity. Since the PCAs of *Nps1/rin^CR^* fruits are in close proximity, hence these were encircled by a single oval. The PCA was constructed using MetaboAnalyst 5.0. The variance of the PC1, PC2, and PC3 are given within parentheses.

**Figure S18.**
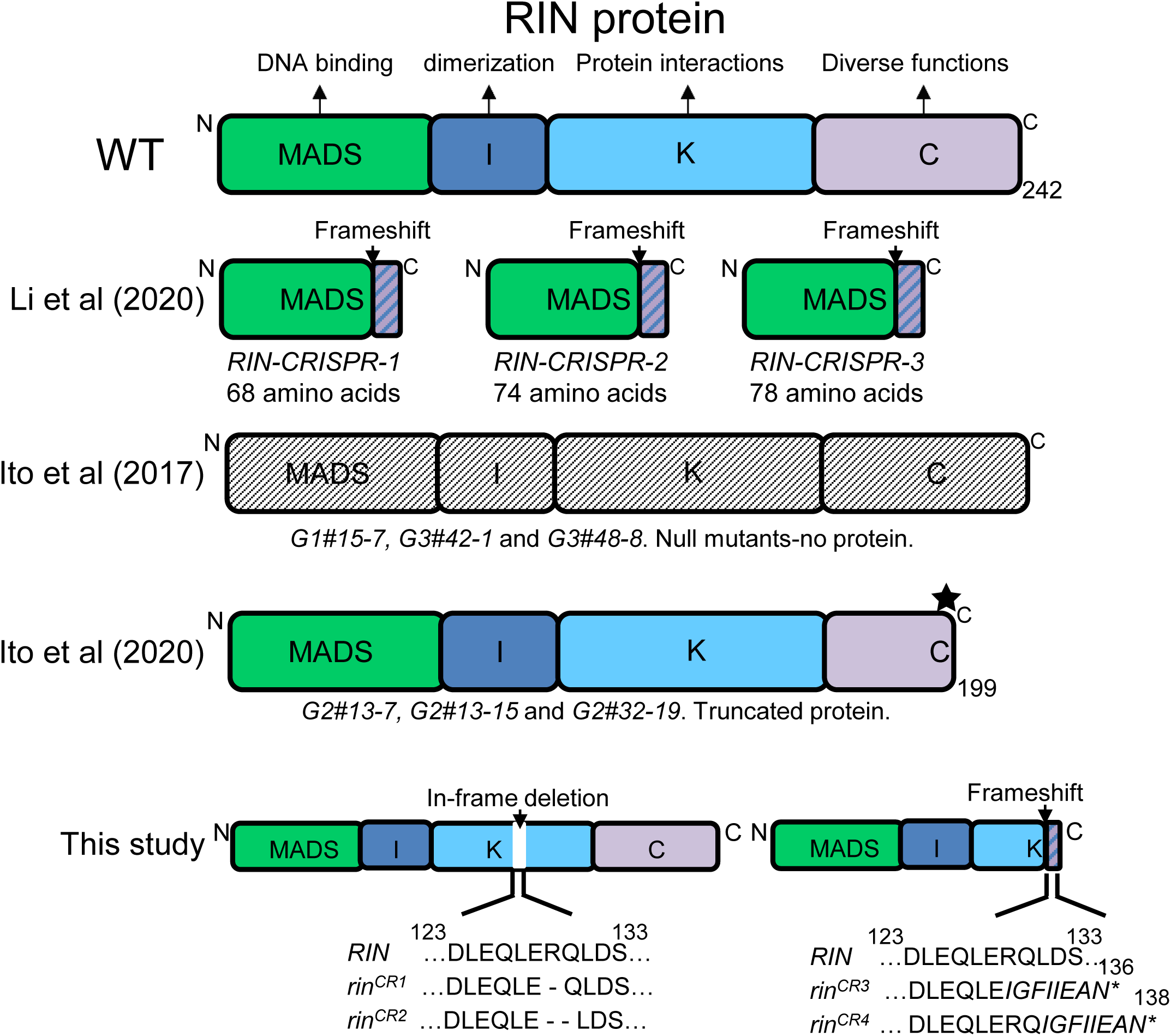
A comparison of genome-edited *RIN* alleles generated in different studies with the present study. The alleles generated by Li, et al. (2020) had frameshift mutations in MADs domain, leading to a truncated protein. The alleles generated by Ito et al. (2017) were likely null alleles, as immunoblots did not detect RIN-specific protein. The rinG2 alleles generated by Ito et al. (2020) encoded a truncated protein. Note the *rin^CR3^* mutant generated in this study is identical to *G1#15-7* (*rinG1-7*) allele generated by Ito et al. (2017). **Ito Y et al. (2017)** Re-evaluation of the rin mutation and the role of RIN in the induction of tomato ripening. *Nature Plants* **3:** 866–874 **Ito Y et al. (2020)** Allelic Mutations in the Ripening -Inhibitor Locus Generate Extensive Variation in Tomato Ripening. *Plant Physiol*. **183:** 80–95 **Li S et al. (2020)** Roles of RIN and ethylene in tomato fruit ripening and ripening-associated traits. *New Phytologist.* **226**: 460-75.

**Figure S19:**
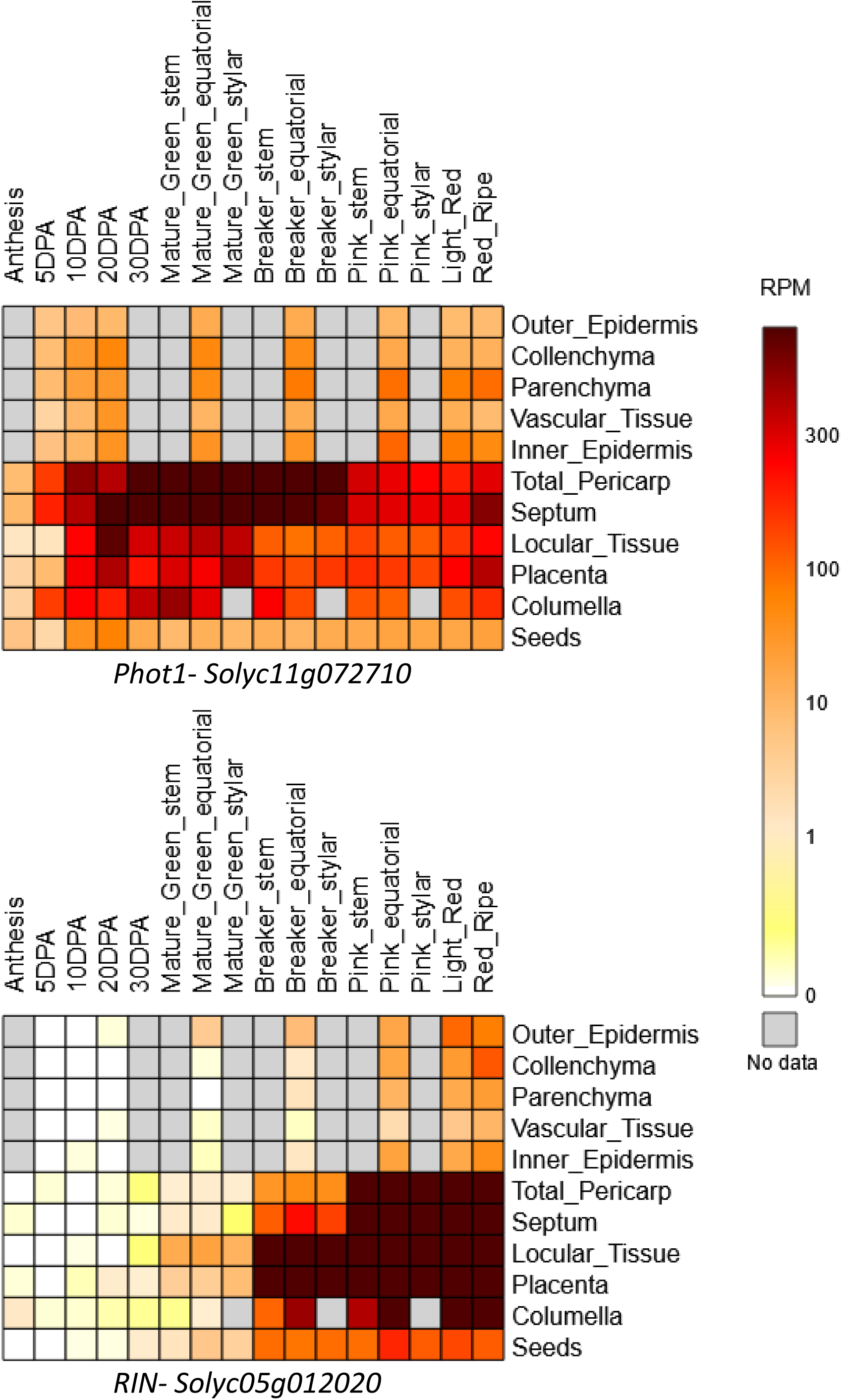
The expression of *RIN* and *phototropin1* genes in tomato fruits. The heat maps show the expression of the above genes in tissues of tomato fruits from anthesis to full ripening (Data source https://tea.solgenomics.net/expression_viewer/output). The side panel shows the colors used in the heat map for different rpm values. Unlike *RIN*, which is a ripening-specific gene, the *phototropin1* gene expresses throughout the fruit development, and during ripening, its expression substantially overlaps with *RIN*.

**Table S1:**
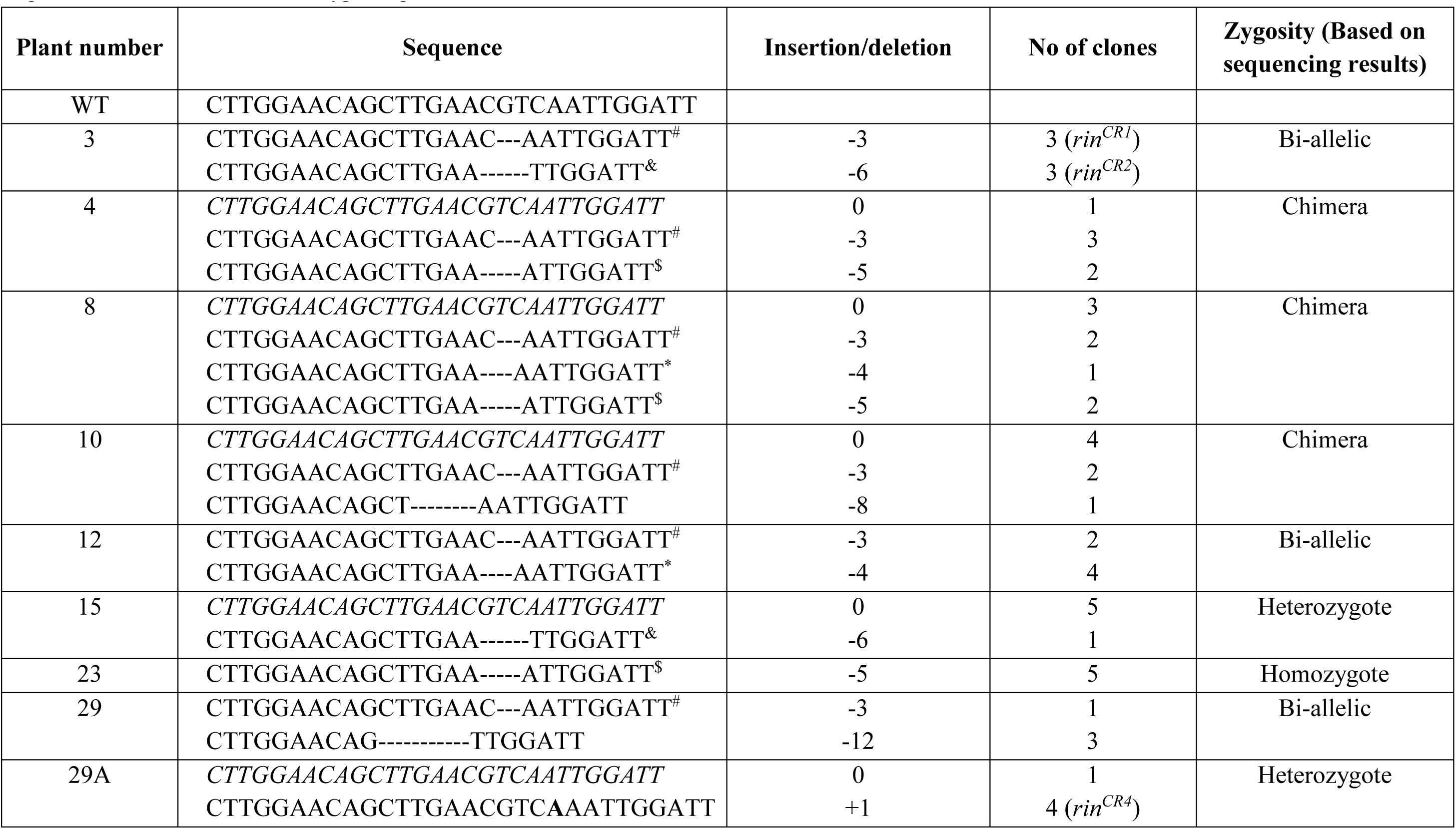
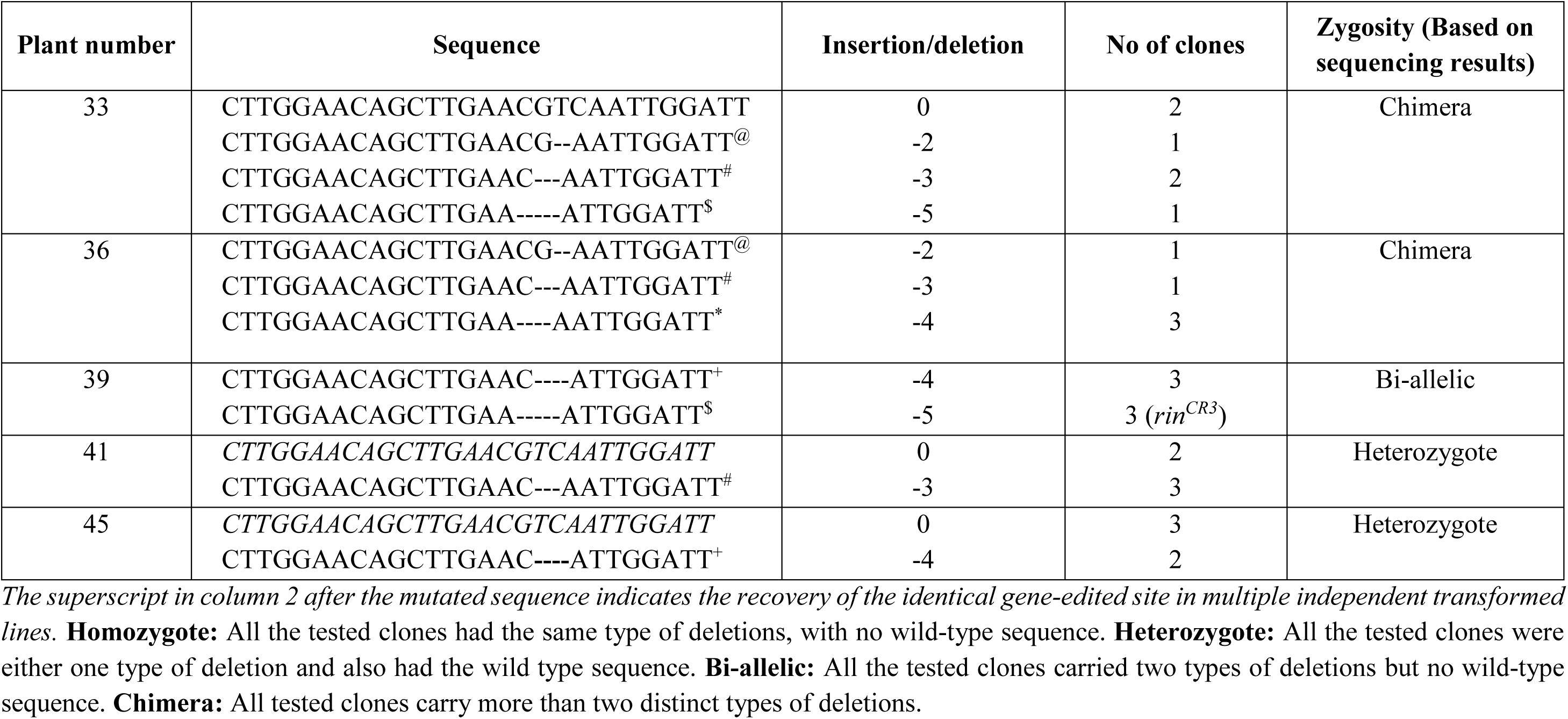
Identification of mutated alleles and zygosity in T_0_ transgenic plants. From individual transgenic plants, the gRNA target region was PCR amplified and cloned into the TA cloning vector. About 6-8 clones were Sanger sequenced. The clones were grouped based on the sequence. The clones with wild-type sequences are marked in italics.

**Table S2:**
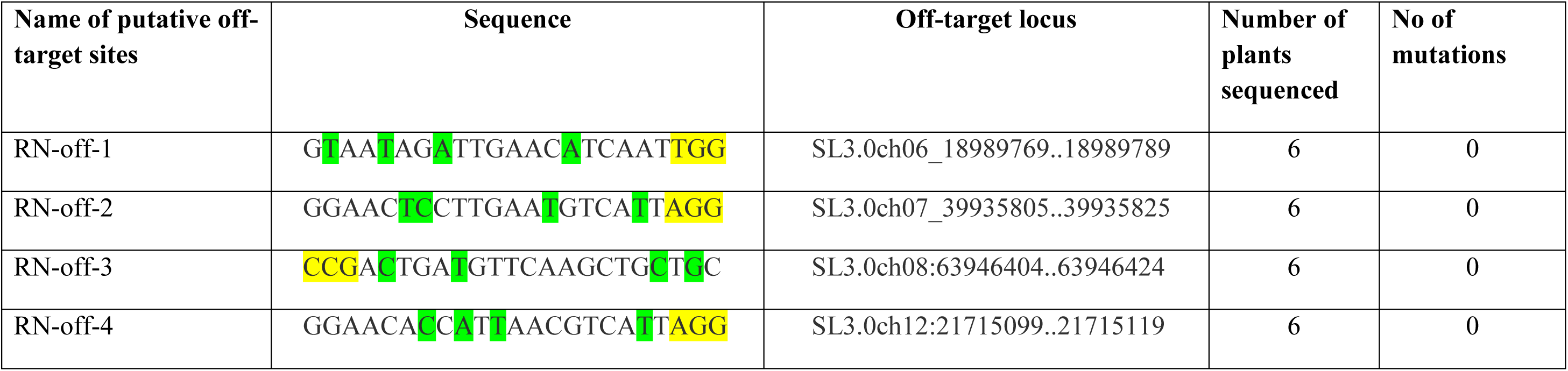
Mutation detection in potential off-target sites. The mismatched nucleotides in the potential off-target locus are highlighted in green, and the PAM sequence (NGG) is highlighted in yellow. Plant numbers 3, 8, 12, 23, 29A, and 39 from the T_0_ generations were examined for off-target mutations. All these plants were Cas9 positive and had confirmed mutagenesis in the target locus.

**Table S3.**
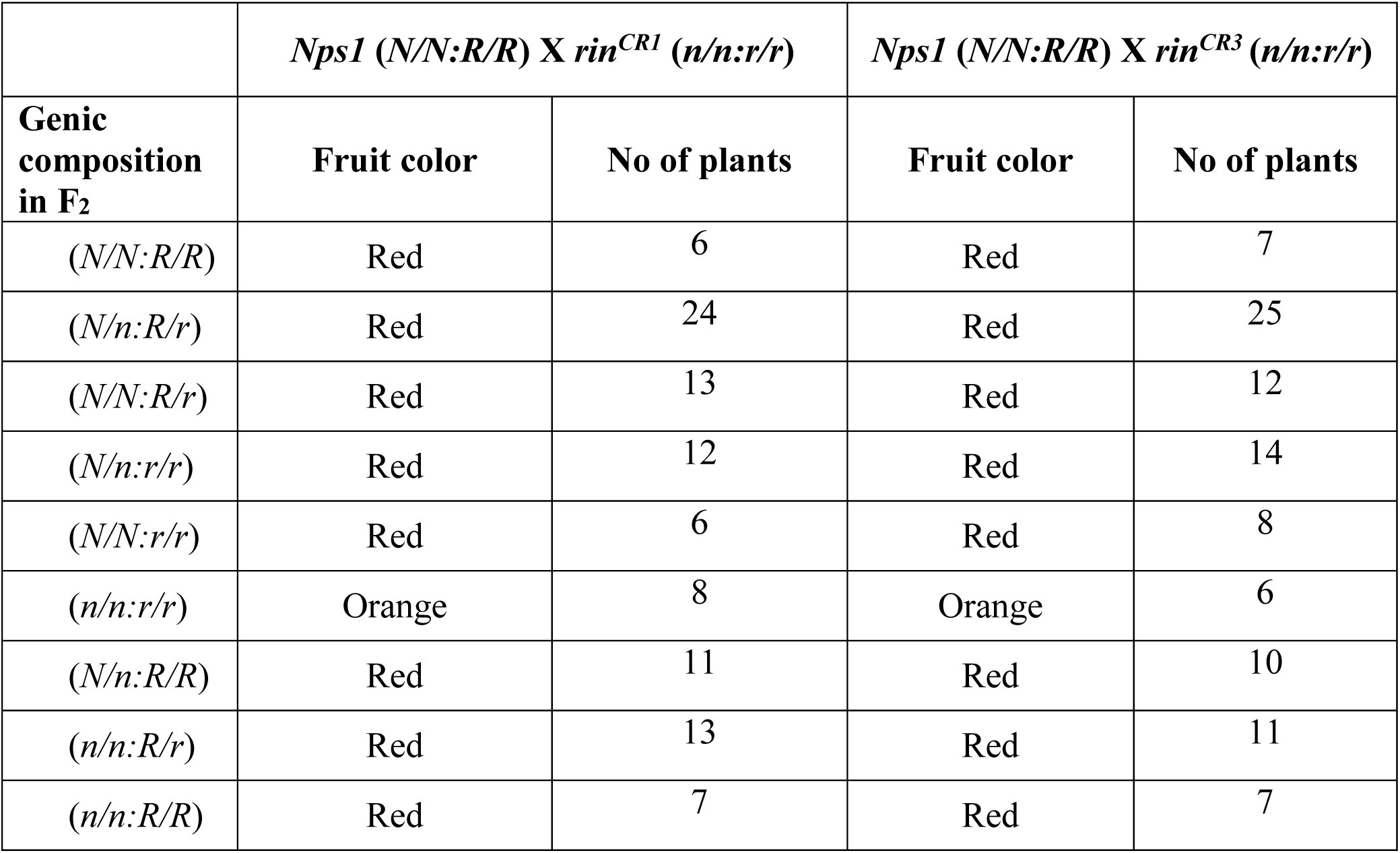
The *Nps1* (*N/N:R/R*) X *rin^CR^* (*n/n:r/r*) (*rin^CR1^* and *rin^CR3^*) crossed F_2_ plants were examined for zygosity of *Nps1* and respective *rin^CR^* mutations at the seedling stage. One hundred genotyped plants were then grown in a greenhouse to monitor the fruit colors. The reappearance of the *rin^CR^* phenotype was observed in plants having the n/n:r/r genotype. Plants were closely monitored for fruit color to distinguish the homozygous *rin^CR^* allele (*n/n:r/r*). The fruits of plants bearing *rin^CR^* allele post-MG stage take much longer to reach the light red color, and late developing fruits on truss stay for a longer duration at the orange stage. For the fruit’s phenotypes, see Figure S10 and Figure S11.

**Table S4.** The fruits of *Nps1* (*N/N:R/R*) X *rin^CR1^* (*n/n:r/r*) crossed F_2_ plants, where both the alleles are present in either homozygous or heterozygous conditions, were monitored for on-vine fruit ripening stages at regular intervals. For fruit phenotypes post RR-stage, see Figure S14. The plants were grown in the greenhouse to monitor the fruit colors.

|  | *Nps1* (*N/N:R/R*) X *rin*<sup>CR1</sup> (*n/n:r/r*) |  |  |  |  |
| --- | --- | --- | --- | --- | --- |
|  | Duration of Fruit ripening stages<br>Days (Mean ± SE) |  |  |  |  |
| Ripening Stages | WT (AV) | (*N/N:r/r*) | (*N/n:R/r*) | (*N/N:R/r*) | (*N/n:r/r*) |
| MG | 36 ± 3 | 48 ± 4 | 47 ± 5 | 48 ± 5 | 48 ± 3 |
| BR | 1 ± 0.04 | 2 ± 0.02 | 1 ± 0.05 | 1 ± 0.02 | 2 ± 0.03 |
| TUR / Orange | 4 ± 0.3 | 4 ± 0.3 | 2 ± 0.04 | 3 ± 0.04 | 6 ± 0.4 |
| Pink | 3 ± 0.1 | 6 ± 0.5 | 4 ± 0.1 | 3 ± 0.02 | 4 ± 0.2 |
| RR | 10 ± 2 | 60 ± 6 | 50 ± 3 | 65 ± 5 | 55 ± 4 |

**Table S5.** The fruits of *Nps1* (*N/N:R/R*) X *rin^CR3^* (*n/n:r/r*) crossed F_2_ plants, where both the alleles are present in either homozygous or heterozygous conditions, were monitored for on-vine fruit ripening stages at regular intervals. For fruit phenotypes post RR-stage, see Figure S15. The plants were grown in the greenhouse to monitor the fruit colors.

|  | *Nps1* (*N/N:R/R*) X *rin*<sup>CR3</sup> (*n/n:r/r*) |  |  |  |  |
| --- | --- | --- | --- | --- | --- |
|  | Duration of Fruit ripening stages<br>Days (Mean ± SE) |  |  |  |  |
| Ripening Stages | WT (AV) | (*N/N:r/r*) | (*N/n:R/r*) | (*N/N:R/r*) | (*N/n:r/r*) |
| MG | 36 ± 3 | 50 ± 3 | 48 ± 4 | 50 ± 6 | 52 ± 5 |
| BR | 1 ± 0.04 | 3 ± 0.2 | 2 ± 0.1 | 2 ± 0.04 | 3 ± 0.05 |
| TUR / Orange | 4 ± 0.3 | 5 ± 0.3 | 4 ± 0.2 | 4 ± 0.3 | 6 ± 0.3 |
| Pink | 3 ± 0.1 | 7 ± 0.6 | 6 ± 0.3 | 6 ± 0.2 | 7 ± 0.3 |
| RR | 10 ± 2 | 75 ± 6 | 55 ± 2 | 70 ± 5 | 77 ± 3 |

**Table S6:**
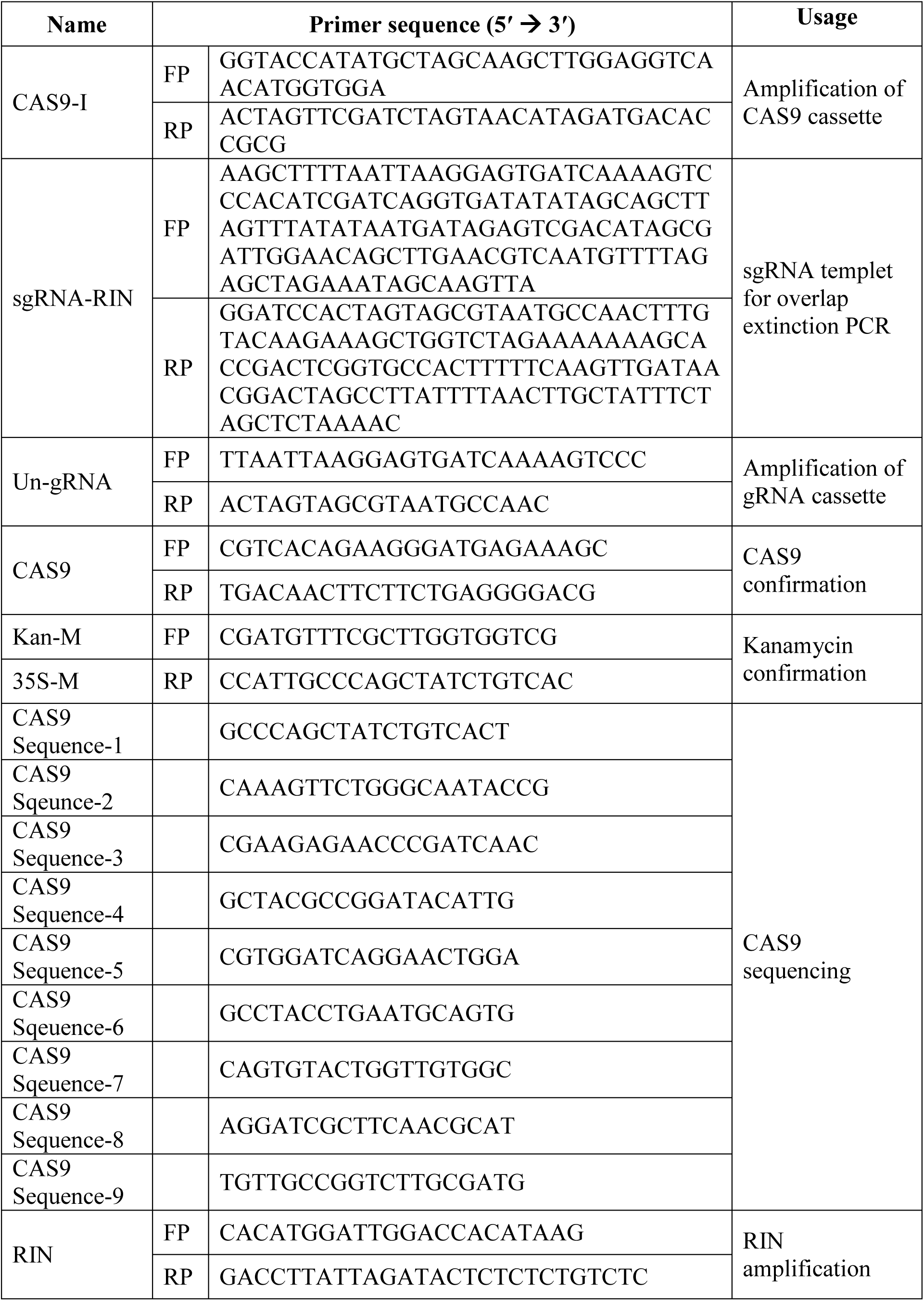
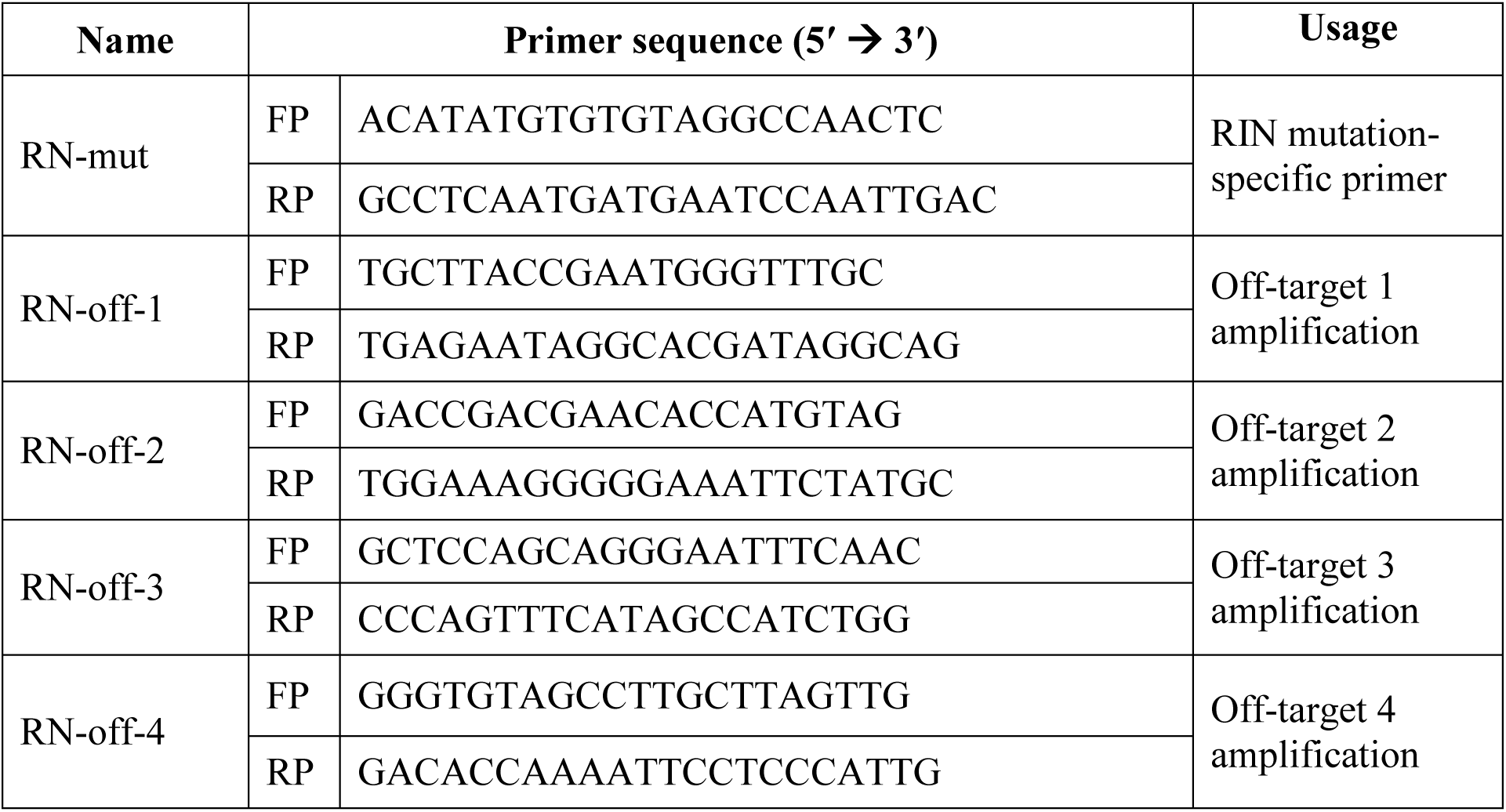
List of primers used for RIN gRNA, plasmid construction, off-target monitoring, PCR-RE assay, and CAS9-free plant monitoring.

**Table S7:**
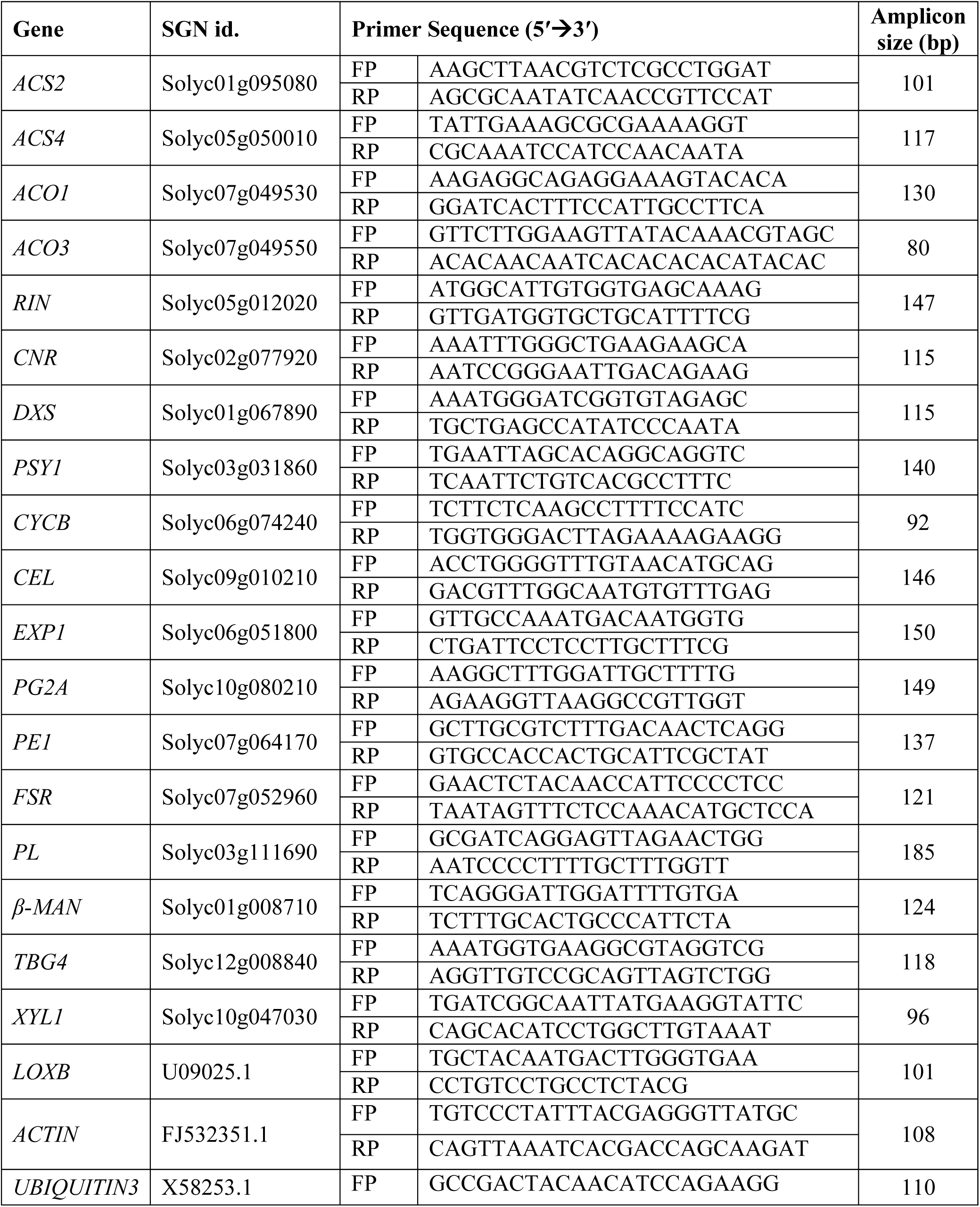
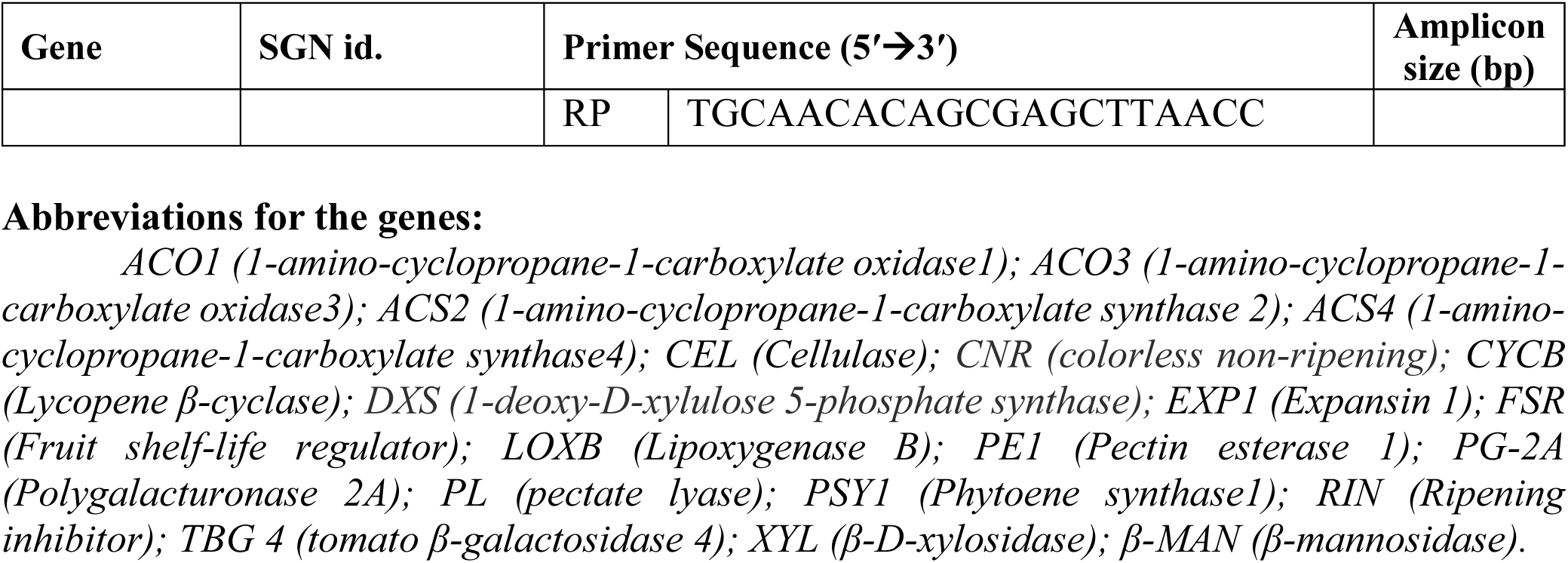
List of primers used for quantitative real-time PCR of genes modulating ethylene biosynthesis, fruit ripening, carotenoids biosynthesis, and cell wall softening. The primers were designed using the SGN SL2.50 version (https://solgenomics.net). *Lipoxygenase B, β-actin, and ubiquitin* sequences were from NCBI (https://www.ncbi.nlm.nih.gov/) unigenes.

